# Co-Dependent and Interdigitated: Dual Quorum Sensing Systems Regulate Conjugative Transfer of the Ti Plasmid and the At Megaplasmid in *Agrobacterium tumefaciens* 15955

**DOI:** 10.1101/2020.12.15.422598

**Authors:** Ian S. Barton, Justin L. Eagan, Priscila A. Nieves-Otero, Ian P. Reynolds, Thomas G. Platt, Clay Fuqua

## Abstract

Members of the *Rhizobiaceae*, often carry multiple secondary replicons in addition to the primary chromosome with compatible *repABC*-based replication systems. Unlike secondary chromosomes and chromids, *repABC*-based megaplasmids and plasmids can undergo copy number fluctuations and are capable of conjugative transfer in response to environmental signals. Several *Agrobacterium tumefaciens* lineages harbor three secondary *repABC*-based replicons, including a secondary chromosome (often linear), the Ti (tumor-inducing) plasmid and the At megaplasmid. The Ti plasmid is required for virulence and encodes a conjugative transfer (*tra*) system that is strictly regulated by a subset of plant-tumor released opines and a well-described acyl-homoserine lactone (AHL)-based quorum-sensing mechanism. At plasmids are generally not required for virulence, but carry genes that enhance rhizosphere survival, and these plasmids are often conjugatively proficient. We report that the At megaplasmid of the octopine-type strain *A. tumefaciens* 15955 encodes a quorum-controlled conjugation system that directly interacts with the paralogous quorum sensing system on the co-resident Ti plasmid. Both the pAt15955 and pTi15955 plasmids carry homologues of a TraI-type AHL synthase, a TraR-type AHL-responsive transcription activator, and a TraM-type anti-activator. The *traI* genes from both pTi15955 and pAt15955 can direct production of the inducing AHL (3-octanoyl-L-homoserine lactone) and together contribute to the overall AHL pool. The TraR protein encoded on each plasmid activates AHL-responsive transcription of target *tra* gene promoters. The pAt15955 TraR can cross-activate *tra* genes on the Ti plasmid as strongly as its cognate *tra* genes, whereas the pTi15955 TraR preferentially biased towards its own *tra* genes. Putative *tra* box elements are located upstream of target promoters, and comparing between plasmids, they are in similar locations and share an inverted repeat structure, but have distinct consensus sequences. The two AHL quorum sensing systems have a combinatorial effect on conjugative transfer of both plasmids. Overall, the interactions described here have implications for the horizontal transfer and evolutionary stability of both plasmids and, in a broad sense, are consistent with other *repABC* systems that often have multiple quorum-sensing controlled secondary replicons.

**CONTRIBUTION TO THE FIELD:** Many bacteria harbor multiple types of plasmids, and in a fraction of these the plasmids encode independent conjugative transfer systems. In the plant pathogen *Agrobacterium tumefaciens*, conjugation of the virulence plasmid, called the Ti (tumor-inducing) plasmid, is regulated by plant released signals called opines, and by a well characterized quorum sensing mechanism also encoded on the plasmid. The co-resident At megaplasmid also has its own conjugative transfer system. In certain lineages of *A. tumefaciens* the At plasmid carries a discrete quorum sensing system. This study aimed to determine the extent to which these plasmid-borne quorum sensing systems overlap to co-regulate conjugative transfer of both plasmids. Co-resident plasmids can impact each other, and the host bacterium at multiple levels, and for the Ti plasmid and At plasmid, we find that each plasmid clearly can influence the conjugative transfer systems of the other plasmid. The findings reported also shed light on how bacteria with multiple quorum sensing pathways have evolved to integrate these systems, in this case to control horizontal gene transfer of two different plasmids. Overall, the interactions described here have broader implications for the horizontal transfer and evolutionary stability of co-resident plasmids, and the degree to which their regulatory systems are coordinated.

## INTRODUCTION

Many members of the *Alphaproteobacteria* (APB) group contain large secondary replicons, in addition to the primary chromosomal replicon with the majority of core genes. These secondary replicons which include megaplasmids, chromids (large replicons, >350 kb, that carry essential genes but are thought to have arisen by horizontal gene transfer), and secondary chromosomes often have *repABC*-based replication origins (Fournes et al. 2018, Pinto et al. 2012). Several APB, notably within the Family *Rhizobiaceae*, harbor multiple large secondary replicons that are also stably maintained by compatible *repABC* systems. Plasmids are common among this family, and a single strain can often harbor multiple plasmids. These plasmids have extensive systems for their retention and proliferation, including copy number control, efficient partitioning with the chromosomes, and conjugative transfer (Pinto, et al. 2012).

The Ti plasmid from plant pathogen *Agrobacterium tumefaciens* is one of the most well-studied *repABC* replicons (Gordon and Christie 2014), in which control of plasmid copy number and horizontal transfer is directly linked to bacterial population dynamics resulting from interactions with host plants. Octopine-type and nopaline-type Ti plasmids of *A. tumefaciens* have been most extensively studied for this regulation, but multiple other agrobacterial and rhizobial plasmids exhibit similar regulatory patterns (He and Fuqua 2006, Wetzel et al. 2015). *A. tumefaciens* causes the plant neoplastic disease crown gall, by driving cross-kingdom horizontal gene transfer to host plants, delivering a segment of the Ti plasmid called the T-DNA, into host cells (Chilton et al. 1977). This process is strictly regulated by plant-released signals via the VirA-VirG two-component system (Winans 1991) which activates virulence (*vir*) gene expression including the VirB Type IV secretion system (T4SS) that delivers the T-DNA to plant cells (Christie 2001). In the plant, the T-DNA is ushered into the nucleus and inserted into the host genome (Gelvin 2003). Stable integration of the T-DNA and expression of the transferred genes results in neoplastic growth of the infected tissue, and visible galls on the infected tissue. These genes also encode proteins which drive production of unusual metabolites called opines that are utilized as semi-exclusive nutrients by agrobacteria in the rhizosphere via catabolic functions primarily encoded on the Ti plasmids (Dessaux et al. 1998). A subset of these opines, called conjugal opines, stimulate interbacterial conjugative transfer of the Ti plasmid, and increased Ti plasmid copy number (Ellis et al. 1982, Klapwijk et al. 1978). In most cases, the conjugal opines act indirectly, by stimulating expression of an acyl-homoserine lactone (AHL) quorum sensing (QS) mechanism. The Ti plasmid encodes a LuxI-type AHL synthase called TraI that synthesizes the AHL *N-*3-oxo-octanoyl-homoserine lactone (3-oxo-C8-HSL) (Fuqua and Winans 1994, Hwang et al. 1994, Moré et al. 1996). As with many other AHL quorum sensing systems, the signal can diffuse across the membrane and out of the cell. However, as the population density of AHL-producing agrobacteria increases, the relative AHL concentration is also elevated, eventually signaling a bacterial quorum. The AHL-dependent, LuxR-type protein TraR, also encoded on the Ti plasmid, interacts with the AHL to activate transcription of conjugal transfer (*tra* and *trb*) genes and the *repABC* replication genes of the Ti plasmid, driving conjugation and increased copy number (Fuqua and Winans 1996, Li and Farrand 2000, Pappas and Winans 2003). In addition, TraR activates *traI* expression, resulting in a positive feedback loop on AHL production. Expression of the Ti plasmid *traR* gene is under the tight transcriptional control of the conjugal opine, through an opine-responsive regulator (Fuqua and Winans, 1994). Therefore, conjugal transfer and copy number of the Ti plasmid is strictly regulated by the plant-tumor-released conjugal opine via *traR* expression, and AHL-producing bacterial population density via the QS mechanism. Other Ti plasmid-encoded functions modulate these processes, most notably the antiactivator TraM which directly inhibits TraR, and maintains the minimum threshold of the protein required to activate *tra*/*trb* and *repABC* expression (Fuqua et al. 1995, Hwang et al. 1995).

In the best studied *A. tumefaciens* strains, conjugal transfer and copy number of the co-resident At megaplasmid is not recognized to be strongly affected by the opine-responsive and quorum sensing control systems which regulate these processes for the Ti plasmid. Rather, At plasmid conjugative transfer is commonly regulated through the *rctA* and *rctB* repressor cascade, which seems to provide largely homeostatic control of At plasmid conjugation (Perez-Mendoza et al. 2005). However, in certain nopaline-type *A. tumefaciens* such as strain C58, the conjugal opines that activate quorum sensing can also stimulate transcription of the *rctA* and *rctB* genes (Lang et al. 2013), thereby providing a regulatory linkage between these plasmids.

Here, we show that the octopine-inducible, quorum sensing-dependent conjugation system of the Ti plasmid from *A. tumefaciens* 15955 interacts directly with the conjugation system of the co-resident At megaplasmid. We find that both plasmids contribute to the same AHL pool through independent *traI* AHL synthases, and that they each also encode AHL-responsive TraR transcription factors that can cross-regulate genes on both replicons, impacting the overall quorum-sensing response as well as conjugative transfer of both the Ti plasmid and the At megaplasmid. The co-regulation of conjugative transfer functions observed here suggests direct coordination between both replicons that may affect their population-level stability and increase overall horizontal transfer. Our observations provide another alternate example of complex interactions between conjugative transfer regulatory elements of co-resident *repABC* plasmids in the rhizobia (Castellani et al. 2019, Cervantes et al. 2019, He et al. 2003). Similar interactions are likely to occur within other systems with multiple *repABC*-based replicons, several examples of which exhibit quorum-sensing controlled replication and horizontal gene transfer systems.

## MATERIALS AND METHODS

### Reagents, media, strains, and growth conditions

All strains, plasmids, and oligonucleotides used in this study are described in Tables S1 and S2. Chemicals, antibiotics, and culture media were obtained from Fisher Scientific (Pittsburgh, PA), Sigma Aldrich (St. Louis, MO), and Goldbio (St. Louis, MO) unless otherwise noted. Plasmid design and verification as well as strain creation was performed as described previously (Morton and Fuqua 2012a) with specific details in the text. Oligonucleotide primers were ordered from Integrated DNA Technologies (Coralville, IA). Single primer extension DNA sequencing was performed by ACGT, Inc. (Wheeling, IL). Plasmids were introduced into *Escherichia coli* via transformation with standard chemically competent preparations and into *A. tumefaciens* via electroporation or conjugation. In-frame deletions of *traI* and *traR* were constructed using a previously described allelic replacement method (Morton and Fuqua 2012a).

*E. coli* was cultured in standard LB broth with or without 1.5% (w v^−1^) agar. Unless noted otherwise, *A. tumefaciens* strains were grown on AT minimal medium containing 0.5% (w v^−1^) glucose and 15 mM ammonium sulfate without added FeS0_4_ (ATGN) (Morton and Fuqua 2012b) or AT with 5% (w v^−1^) sucrose as the sole carbon source (ATSN). When required, appropriate antibiotics were added to the medium as follows: for *E. coli*, 100 μg ml^−1^ ampicillin (Ap), 50 μg ml^−1^ gentamicin (Gm), and 50 μg ml^−1^ kanamycin (Km); and for *A. tumefaciens*, 300 μg ml^−1^ Gm, 300 μg ml^−1^ Km, and 300 μg ml^−1^ spectinomycin (Sp). When indicated, *A. tumefaciens* 15955 strains were grown in ATGN media supplemented with 3.25 mM octopine (Octopine, + Oct.) and/or 0.1 μM synthetic 3-oxo-octanoyl-L-homoserine lactone (+AHL, 3-oxo-C8-HSL).

### Genome sequence and bioinformatic analysis

The genome assembly for *A. tumefaciens* 15955 was deposited into GenBank by Academia Sinica (Genbank Access. No. 7601828). Multiple and pairwise sequence alignments were performed with Clustal Omega (Sievers et al. 2011) using default parameters. Promoter sequence motifs were generated with WebLogo (Crooks et al. 2004).

### Controlled expression and reporter plasmids

Plasmid-borne expression constructs were created using the LacIQ encoding, IPTG-inducible series of pSRK vectors (Khan et al. 2008). Coding sequences of *traR* and *traI* from pAt15955 (At15955_50350 and At15955_49060, respectively) and pTi15955 (At15955_53990 and At15955_54620, respectively) were PCR amplified from *A. tumefaciens* 15955 gDNA using primers for the corresponding gene (Table S3) and Phusion DNA polymerase (New England Biolabs, Beverly, MA). PCR products were ligated into pGEM-T easy (Promega, Madison, WI), excised by restriction enzyme digest, and ligated into either pSRKKm or pSRKGm. Final pSRK constructs were confirmed with sequencing and restriction enzyme digestion prior to electroporation into *A. tumefaciens*.

Promoter expression reporter plasmids were created in pRA301 (Akakura and Winans 2002a). Promoter regions for *traA* (At15955_53920 for pTi15955 and At15955_50400 for pAt15955) and *traI* (~500 bp) were amplified with PCR (as above) using *A. tumefaciens* 15955 gDNA and the appropriate primers (Table S3). All reporter gene fusions have *lacZ* translationally fused in frame at the start codon of each gene. PCR products were then assembled into restriction enzyme digests of pRA301 in a two-part NEBuilder isothermal assembly reaction (New England Biolabs, Beverly, MA), verified by PCR and sequencing, and introduced into *A. tumefaciens* via electroporation.

### Promoter expression assays

For octopine and AHL induction of *traA* and *traI* promoters, *A. tumefaciens* 15955 carrying the appropriate pRA301-based reporter constructs were grown to an optical density at 600 nm (OD_600_) of 0.4-0.6 in ATGN supplemented with appropriate antibiotics and spotted onto 0.2 μm sterile filter disks (0.45 μm pore size) placed on small ATGN plates (35 x 10 mm) supplemented with octopine (3.25 mM) and/or AHL (0.1 μM 3-oxo-C8-HSL) and incubated for 48 hours at 28°C. *A. tumefaciens* 15955 strains were then removed from the filter discs, diluted to an OD_600_ ~0.6 in sterile water and used as in standard β-galactosidase assays, as previously described (Miller 1972).

To determine the effect of *traR* expression on promoter activities of *P_traA_* and *P_traI_* from pAt15955 and pTi15955, A*. tumefaciens* 15955 and *A. tumefaciens* NTL4 derivatives carrying pSRK-based *traR* expression constructs and pRA301-based *lacZ* reporter constructs (Table S1) were grown in ATGN liquid cultures with appropriate antibiotics and 400 μM IPTG in the presence or absence of AHL to an OD_600_ ~0.6 and used in standard β-galactosidase assays.

### AHL detection assays

For AHL detection on solid media, we utilized the AHL biosensor strain [*A. tumefaciens* NTL4 (pCF218)(pCF372)] (Fuqua and Winans 1996). This biosensor relies on a high expression, plasmid-borne copy of the AHL-responsive TraR protein from *A. tumefaciens* R10 (pCF218), and a second plasmid carrying a TraR-dependent promoter fused to *lacZ* (*traI-lacZ*, pCF372), both harbored in *A. tumefaciens* NTL4 Ti-plasmidless strain which lacks an AHL synthase, and therefore does not produce endogenous AHLs. The biosensor is a sensitive detection system for exogenous AHLs, expressing *lacZ* only when inducing AHLs are present (McLean et al. 1997). Strain derivatives to be tested for AHLs were grown overnight to an OD_600_ 0.4-0.6 in ATGN supplemented with appropriate antibiotics and then each strain to be tested was struck onto ATGN agar supplemented with 5-bromo-4-chloro-3-indolyl-β-D-galactopyranoside (X-Gal) ~1 cm from the AHL biosensor strain. Media used in assays to test for AHL production with *A. tumefaciens* NTL4-derivatives also had 400 μM IPTG unless otherwise noted. The AHL biosensor was also used to detect AHL in cell-free supernatant from *A. tumefaciens* strains. Strains to be tested were grown overnight in ATGN supplemented with appropriate antibiotics to an OD_600_ of 0.4-0.6 and 1 mL of culture was then pelleted, supernatant removed and re-centrifuged to remove additional cell debris, and filter sterilized (0.2 μm) to create cell-free supernatant (CFS). 100 μL of CFS was then added to cultures of the AHL biosensor, grown to an OD_600_ 0.4-0.6, and assayed for promoter activity using standard β-galactosidase assays. Relative AHL levels for CFS were then reported as Miller units normalized to the OD_600_ of the tested culture.

### Conjugation experiments

Conjugation assays were performed as described previously (Heckel et al. 2014) with alterations. Briefly, liquid cultures of donors and recipients were grown to mid-logarithmic phase in ATGN at 30°C and normalized to OD_600_ ~0.6, mixed 1:1, and 100 μL was spotted onto 0.2 μm pore size cellulose acetate filter disks placed on small ATGN plates (35 x 10 mm). Conjugations were performed on ATGN with *A. tumefaciens* 15955 ectopically expressing plasmid-borne *P_lac_-traR* fusions with or without 400 μM IPTG and appropriate antibiotics. For AHL rescue of conjugation in the *traI* double mutant, varying amounts of 3-oxo-C8-HSL were also added. Following incubation (18h, 28°C), cells were suspended in liquid ATGN, serially diluted, and spotted onto media supplemented with appropriate antibiotics to enumerate donors and transconjugants. Conjugation efficiencies were determined as the ratio of transconjugants per output donor.

#### Statistical analysis

Assays were performed with at least three biological replicates. Standard deviation was determined and presented as error bars. Where appropriate, significance was conservatively evaluated using 2-sided, paired *t*-tests.

## RESULTS

### Conjugation systems of pAt15955 and pTi15955 are activated by octopine

*A. tumefaciens* 15955 is a member of the Genomospecies Group 1 along with similar derivatives *A. tumefaciens* Ach5 and A6 (Lassalle et al. 2011). Sequence analysis of pAt15955 identified conjugation genes homologous to the quorum-sensing regulated *tra*/*trb* system encoded on pTi15955 and other Ti plasmids (Alt-Mörbe et al. 1996, Farrand et al. 1996), including *tra* and *trb* structural genes and homologues of the quorum sensing regulators *traI, traR*, and *traM* (Figure S1A). The conjugation system encoded on pAt15955 is distinct from those found on many other At plasmids, including those of the type strain *A. tumefaciens* C58, a nopaline-type strain (Genomospecies 8), in which pAtC58 conjugates via a *virB-*like type IV secretion system (*avhB*) (Chen et al. 2002) regulated by *rctA/rctB*-like regulators commonly found for other rhizobial plasmids (Nogales et al. 2013, Perez-Mendoza, et al. 2005) (Figure S1). Given the presence of TraI, TraR and TraM homologs on pAt15955 (Figure S1B and S1C), it was conceivable that the conjugative transfer of this plasmid might be regulated by quorum sensing and that pAt15955 and pTi15955 could contribute and respond to a common AHL pool. We previously showed that pAt15955 encodes a functional conjugation system and observed that conjugation of pAt15955Δ270 (a derivative with a 270 kb deletion on pAt) which lacks *cis*-encoded *trb* conjugation functions (Figure S1A), could be stimulated by octopine-induction in a pTi15955-dependent manner (Barton et al. 2019) suggesting a direct interaction between the conjugation systems of both plasmids.

To compare the impact of octopine and AHL cues on the expression of pAt15955 and pTi15955 conjugative transfer (*tra*) genes, we constructed independent translational *lacZ* fusions to the upstream regions and start codons of the predicted *traA* nickase/helicase homologues and the *traI* AHL synthase homologues, carried on the broad host range plasmid pRA301 (Akakura and Winans 2002b), Table S2, Methods). These fusion plasmids were introduced into wild type *A. tumefaciens* 15955, and used in β-galactosidase assays to measure gene expression in the presence or absence of 0.1 μM 3-oxo-C8-HSL and/or 3.25 mM octopine (Figure 1, Materials and Methods). The β-galatosidase activity imparted by the *traA-lacZ* fusions from pTi15955 (green bars) and pAt15955 (blue bars) was low in the absence of octopine or AHL addition (Figure 1A). Addition of octopine alone (medium-fill bars) or octopine plus AHL (dark-fill bars) significantly increased activity from the *traA*^At^-*lacZ* and *traA*^Ti^-*lacZ* fusions, although overall activity of each was relatively low. The same pattern of regulation was observed for 15955 harboring the *traI-lacZ* fusions, although the level of activity from both *traI*^Ti^-*lacZ* fusions was considerably higher than the *traA-lacZ* fusions, and addition of octopine and AHL together had a stronger effect than octopine alone (Figure 1B). The *traI*^At^-*lacZ* fusion (blue bars) showed a similar pattern although its overall expression was lower than the *traI*^Ti^ fusion (green bars). Taken together with our prior observation of pTi15955-dependent octopine-induction of pAt15955 conjugation (Barton, et al. 2019), this data suggests that pTi15955 is directly involved in activation of pAt15955 *tra* gene expression in response to octopine (Fuqua and Winans 1996, Fuqua and Winans 1994). Additionally, octopine-induction had differential effects on pAt15955 and pTi15955 gene activation, where octopine addition stimulated comparable modest expression from both *traA* genes (Figure 1A), but had a more dramatic effect on *traI*^Ti^ expression than that of *traI*^At^ (Figure 1B), suggesting complex cross-regulation between both plasmids and supporting our previous observations for pAt15955 conjugation control, as well as previous work on the activation of octopine-type Ti plasmid conjugation (Fuqua and Winans 1996, Fuqua and Winans 1994).

**Figure 1.**
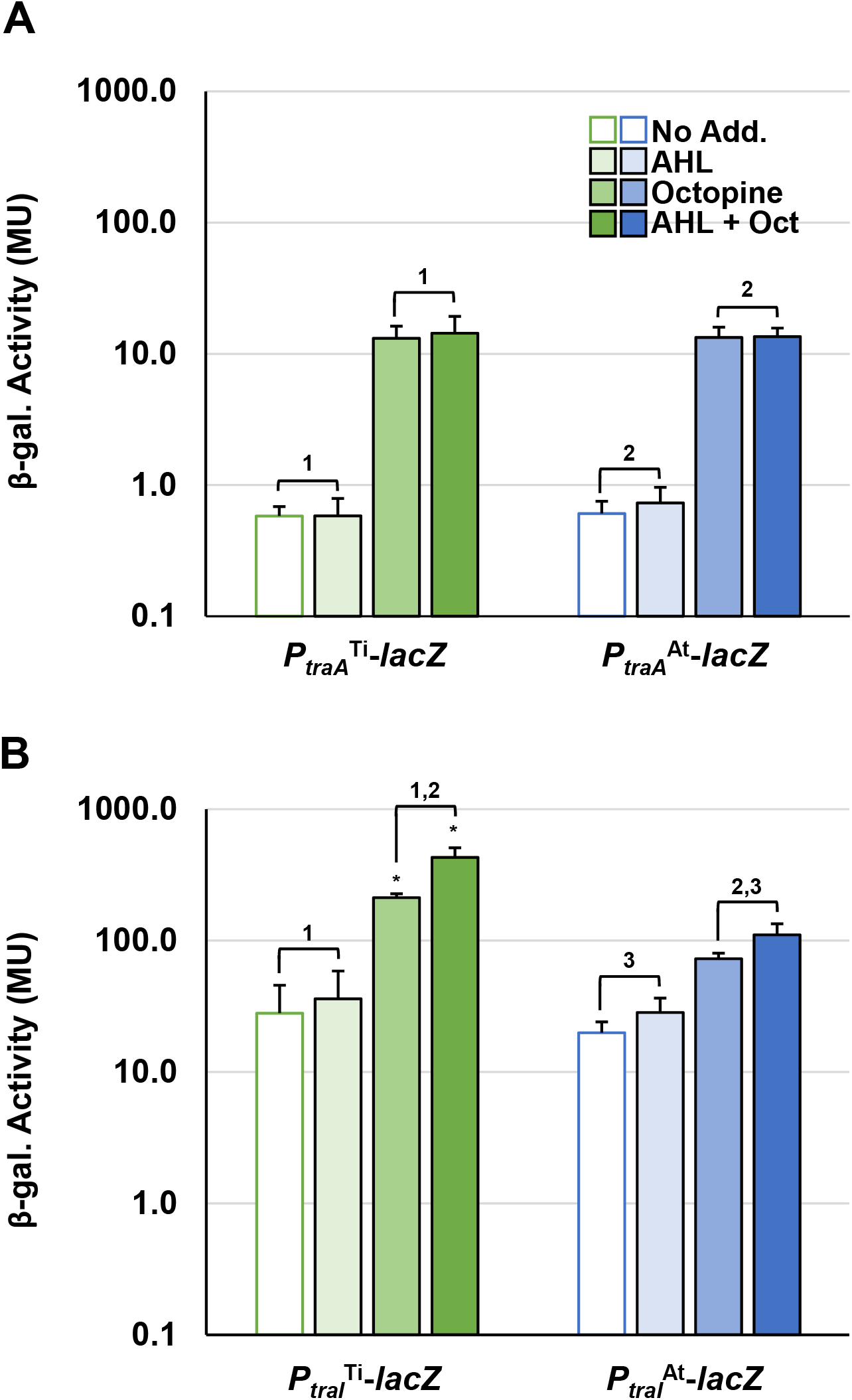
Octopine-induced activation of *P_traA_* and *P_traI_* from pAt and pTi in WT *A. tumefaciens* 15955. WT *A. tumefaciens* 15955 carrying a plasmid-borne copy of either *P_traA_* (top) or *P_traI_* (bottom) from pTi15955 (green) or pAt15955 (blue) translationally fused at their start codons to *lacZ* were spotted on 0.2 μm cellulose acetate filter discs placed on ATGN with or without 3.25 mM octopine (Oct.) and/or 0.1 μM N-3-oxo-C8-HSL (AHL) and incubated at 28°C for 48 hours. Bars with no, light, medium, or dark fill indicate no addition, addition of AHL, addition of octpine, or addition of both octopine and AHL, respectively. Cells were resuspended and promoter activities were calculated as Miller Units (Methods). Bars under the same bracket are not significantly different, unless they are marked with an asterisk (p-value <0.05). Bars under brackets with that have the same number above them (1, 2, 3) in each panel (A/B) denote a significant difference in averages (p-value <0.05) between the bars under the similarly numbered brackets.

### *A. tumefaciens* 15955 encodes two functional homologs of the *traI* AHL synthase

The pAt15955 plasmid has two annotated *traI* homologs, one of which is encoded as the predicted first gene of a *traI-trb* operon (*traI*^At_1^, At15955_49060) in a position similar to pTi-encoded systems, and another ~20 Kbp distant from *trb* in an uncharacterized locus (*traI*^At_2^, At15955_RS25205) (Figure S1A and S1C). Both are within the large segment of pAt15955 that is deleted upon curing of pTi15955 (purple bar in Figure S1), and a pAt15955 derivative with this deletion is unable to conjugate independently (Barton et al. 2019). The annotated coding sequence of *traI*^At_2^ specifies a protein that has a 25 amino acid C-terminal extension relative to TraI^Ti^ and TraI^At_1^ (Figure S2). TraI^At_1^ is 100% identical to a TraI homologue we identified earlier from the related Genomospecies 1 strain *A. tumefaciens* A6, that we found to specify production of 3-oxo-C8-HSL (Wang et al. 2014). To evaluate the functionality of the three *traI* homologs in *A. tumefaciens* 15955, each ORF was fused in-frame with the *lacZ* start codon under control of *P_lac_* in the broad host range vector, pSRKGm, to create the following constructs: pSRKGm *P_lac_*-*traI*^At_1^ (pIB302), pSRKGm *P_lac_*-*traI*^At_2^ (pIB305) and pSRKGm *P_lac_*-*traI*^Ti^ (pIB303) (Table S2). These plasmids were introduced individually into *A. tumefaciens* NTL4, a strain that lacks the Ti plasmid and does not produce AHL (Table S1) to create derivatives that were used in agar plate-based AHL crossfeeding assays (McLean, et al. 1997) with the *A. tumefaciens* NTL4(pCF218)(pCF372) AHL biosensor (Figure 2A). With this biosensor, diffusible AHLs in the producing strain cause induction of the AHL-responsive biosensor resulting in blue X-Gal pigmentation via increased β*-*galactosidase activity. The biosensor responds optimally to 3-oxo-C8-HSL, but will also detectably respond at diminishing levels to a broad range of non-cognate AHLs, from acyl chain lengths of C6-C14, and either 3-oxo-, 3-OH or fully reduced at the β-carbon (Zhu et al. 1998). Expression of either normal length *traI* from pAt15955 or pTi15955 grown in the presence of 400 μM IPTG to induce expression of the *P_lac_-traI* fusion was able to generate detectable AHL via the plate-based assay, validating that both genes are functional (Figure 2A). However, no AHL production was detected from the strain harboring the *traI*^At_2^ plasmid, suggesting it is nonfunctional or produces a product not recognized by the AHL biosensor strain.

**Figure 2:**
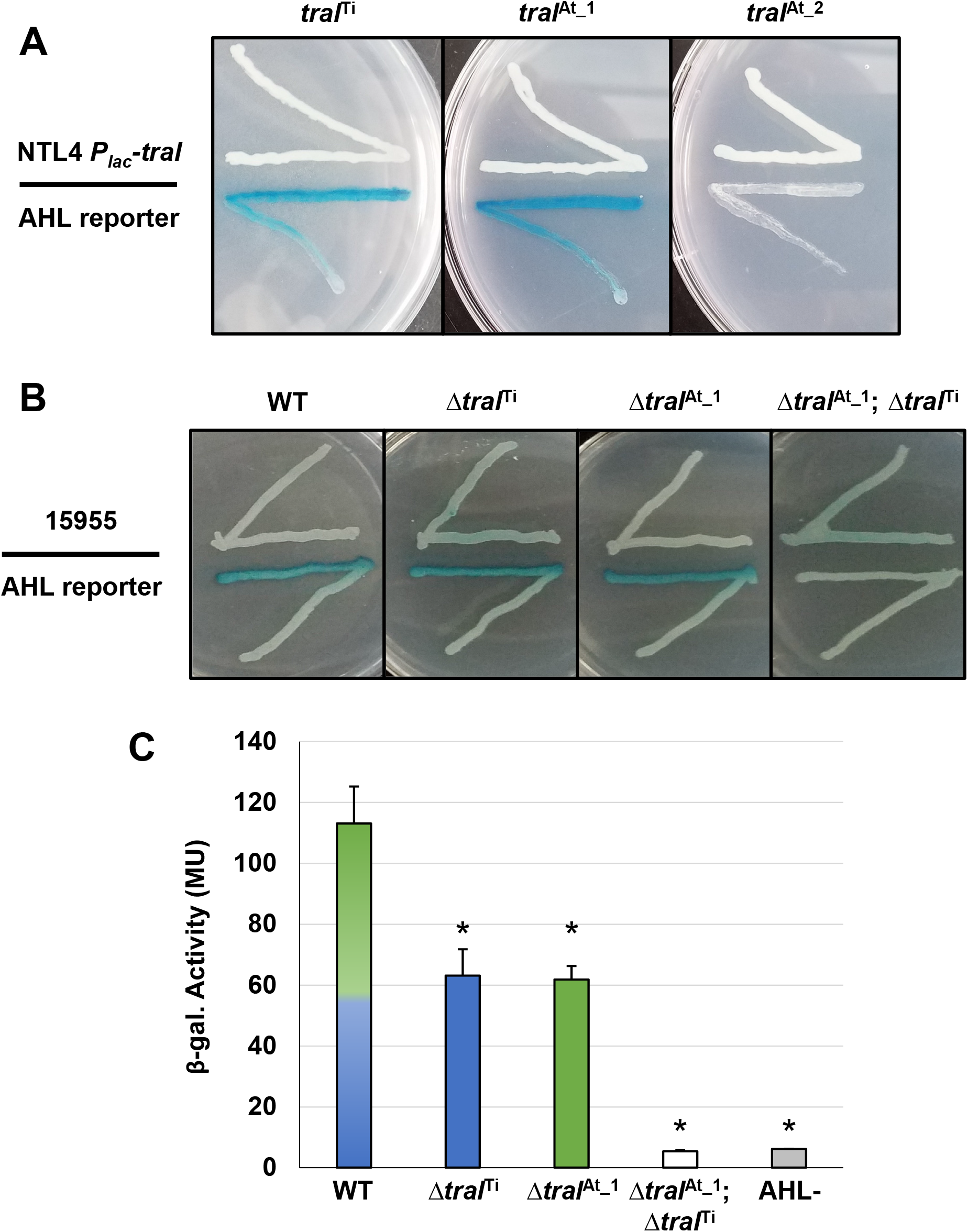
Qualitative and quantitative AHL assays of *A. tumefaciens* strains ectopically expressing *traI* genes, and *traI* mutant derivatives. **(A)** AHL production by *A. tumefaciens* NTL4 (Ti plasmidless, AHL^−^) ectopically expressing *traI* genes from pAt15955 and pTi15955 expressed from *P_lac_* (top streaks). These strains were used in a plate-based AHL production diffusion assays (McLean, et al. 1997) with NTL4(pCF218)(pCF372), (AHL reporter, bottom streak). The *traI* designations are listed above. Media was ATGN agar supplemented with IPTG (400 μM) and X-Gal (40 ug/ml). **(B)** *A. tumefaciens* 15955 or derivatives with clean deletions in *traI* from pTi15955 (Δ*traI*^Ti^), pAt15955 (Δ*traI*^At_1^) or both (Δ*traI*^Ti^; Δ*traI*^At_1^) (top streak) were used in plate-based AHL production assays with *A. tumefaciens* NTL4(pCF218)(pCF372) (AHL reporter, bottom streak). **(C)** Cell-free supernatants from *A. tumefaciens* 15955 (WT, mixed green/blue bar) and derivatives mutated for *traI* on pTi15955 (Δ*traI*^Ti^, blue bar, pAt15955 (Δ*traI*^At_1^, green bar), or both (Δ*traI*^Ti^; Δ*traI*^At_1^, open bar) were added to subcultures of the AHL biosensor strain, *A. tumefaciens* NTL4(pCF218)(pCF372), and allowed to grow until mid-exponential phase. Standard β-gal assays were then conducted on reporter strain cultures to determine relative AHL levels in the cell-free supernatant, calculated as β-gal activity (Methods). AHL-indicates cell-free supernatant from the biosensor strain *A. tumefaciens* NTL4(pCF218)(pCF372) (grey bar). Assays performed as three biological replicates, and bars marked with asterisks are significantly different from wild type (p-value <0.05).

To determine whether both functional *traI* genes are expressed in WT *A. tumefaciens* 15955 and contribute to the overall AHL pool, plate-based AHL detection assays were conducted (as above) with WT *A. tumefaciens* 15955 and strains isogenic for deletion of *traI* gene from the Ti plasmid (Δ*traI*^Ti^), the At plasmid (Δ*traI^At_1^*), or both (Δ*traI*^Ti^Δ*traI^At_1^*) (Table S1, Figure 2B). AHL production was detected for WT *A. tumefaciens* 15955 and the individual deletion mutants, but the double mutant did not produce inducing levels of AHL (Figure 2B), indicating that both *traI* genes are expressed in *A. tumefaciens* 15955 and each contribute to detectable levels of AHL. To assess the relative amount of AHL produced by each TraI protein in standard growth media, quantitative β-gal assays were conducted with the AHL reporter (NTL4 pCF218 pCF372) using cell-free supernatant from the isogenic *traI* mutant strains (Figure 2C). Cell-free supernatant from turbid cultures from either *traI* mutant strain provided half of the B-gal activity relative to WT 15955, which indicates roughly equal contribution from each *traI* under standard growth conditions in the absence of octopine. The double deletion mutant resulted in background activity equivalent to an AHL-control strain (Figure 2C).

### *traI*-dependent activation of pAt15955 and pTi15955 conjugation by *traR*^Ti^

One explanation for octopine-induced activation of pAt15955 conjugation is direct regulation of pAt15955 *tra* genes by *traR*^Ti^. To determine the effect of *traR*^Ti^ on conjugation in *A. tumefaciens* 15955, we generated *P*_lac_-*traR* constructs fused in-frame at the *lacZ* start codon and carried on pSRKKm (pIB309) or pSRKGm (pIB307). These plasmids were independently introduced into WT *A. tumefaciens* 15955 derivatives that carry antibiotic resistance marked derivatives of either pTi15955 (IB123, Gm^R^) or pAt15955 (IB125, Km^R^), respectively, and assayed for conjugation efficiencies of pTi15955 or pAt15955 upon induction of *traR*^Ti^ with 400 μM IPTG (Figure 3, green and blue bars, respectively). As with other Ti plasmids, pTi1955 plasmid does not conjugate at detectable levels in the absence of octopine, but this requirement can be bypassed with ectopic expression of *traR* (Fuqua and Winans 1994). Expression of *traR*^Ti^ stimulated conjugation of pTi15955 to a level consistent with our previous findings, and also strongly stimulated pAt15955 conjugation (Figure 3).

**Figure 3:**
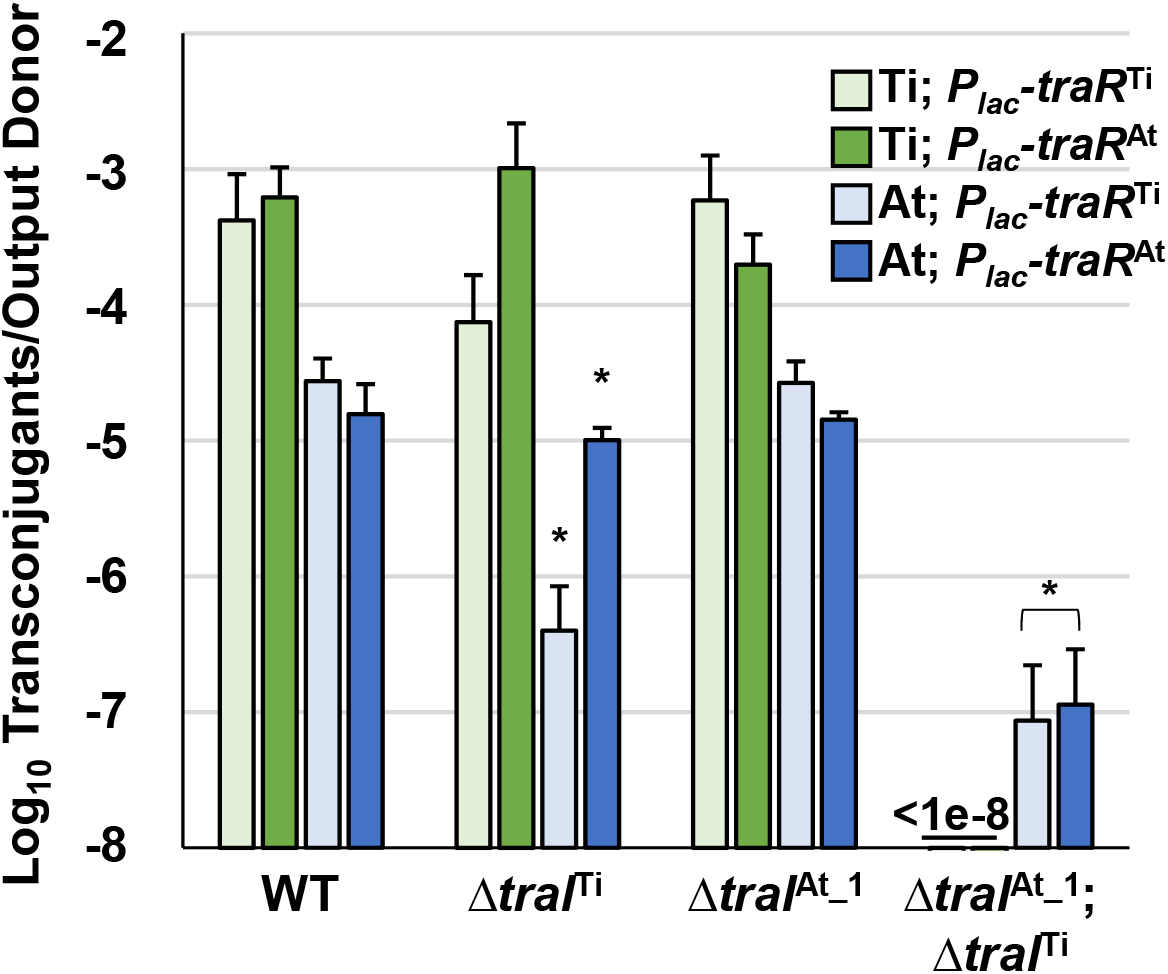
Stimulation of pTi15955 and pAt15955 conjugation by *traR* expression is dependent on pAt15955- and pTi15955-encoded *traI*. *A tumefaciens* 15955 derivatives with a GmR cassette integrated at a neutral location on pTi15955 (green bars) or a KmR cassette similarly integrated on pAt15955 (blue bars) harboring either a plasmid-borne copy of *P_lac_*-*traR* from pTi15955 (light green and light blue bars) or pAt15955 (dark green and dark blue bars) were mixed 1:1 with a plasmidless recipient (*A. tumefaciens* ERM52, SpR) and spotted on ATGN agar containing 400 μM IPTG. After 24 hours incubation, transconjugants and donors were enumerated and conjugation frequencies were calculated as transconjugants per output donor (Methods). Green and blue bars depict pTi15955 and pAt15955 conjugation frequencies, respectively. Asterisks denote significant difference between averages relative to wild type (p-value <0.05).

As with many quorum sensing systems, stimulation of conjugation by *traR*^Ti^ in previously studied *A. tumefaciens* strains requires elevated levels of AHL, augmented by positive feedback on AHL production, to fully activate TraR by stabilizing functional dimerization and DNA binding (Hwang et al. 1999, Qin et al. 2000). To test how the two *traI* genes contribute to AHL-mediated induction of pTi15955 and pAt15955 conjugation, experiments were performed as above in derivatives deleted for *traI*^Ti^ *or traI*^At_1^, or both. Interestingly, conjugation of pTi15955 and of pAt15955 was activated by expression of *traR*^Ti^ in the presence of either version of *traI*, but was extremely low or undetectable (<10^−7^ conjugation efficiency) when both were mutated (Figure 3 light green and light blue bars). However, while expression of *traR*^Ti^ was able to stimulate equivalent levels of conjugation in the wild type or in the Δ*traI*^At_1^ mutant, deletion of the *traI*^Ti^ decreased conjugation frequency of both plasmids by 10-fold or greater, although it was still detectable over the Δ*traI*^Ti^Δ*traI*^At_1^ double mutant. Conjugation of pAt15955 from the Δ*traI*^Ti^Δ*traI^At^* mutant could be rescued with exogenous addition of 3-oxo-C8-HSL, confirming that the extremely low conjugation from this mutant was due to differences in AHL levels (Figure S3).

### *traR*^At^ is a potent activator of pAt15955 and pTi15955 conjugation

LuxR-type proteins are defined by several well conserved amino acid positions in the N-terminal domain that mediate AHL recognition, and in their C-terminal domains that usually coordinate DNA binding. TraR^At^ is very similar to TraR^Ti^ (63.2% identity) and both share the majority of these defining conserved residues (Figure S4). However, they differ at position 194 where TraR^Ti^ has a well conserved isoleucine (I194) residue whereas at this position TraR^At^ has an alanine (A194) residue, an amino acid residue not found in most other LuxR homologues (Whitehead et al. 2001). To determine whether the *traR* homologue from pAt15955 could stimulate conjugation of either pAt15955 or pTi15955, conjugation experiments were performed as above with ectopic expression of *P_lac_*-*traR*^At^ (Figure 3, dark green and dark blue bars). The *traR*^At^ plasmid was able to stimulate conjugation of pTi15955 and pAt15955 to a magnitude similar for *traR*^Ti^ in a *traI*-dependent manner (Figure 3). However, in contrast to *traR*^Ti^, *traR*^At^ was able to effectively stimulate conjugation of pTi15955 and pAt15955 when either *traI* gene, *traI*^Ti^ or *traI*^At_1^ was present (Figure 3, dark green and dark blue bars), suggesting that *traR*^At^ can potentiate AHL production from either *traI*.

### Differential activation of AHL production by TraR^Ti^ and TraR^At^ depends on each *tral* gene

Our findings have revealed that the *traI* genes from pAt15955 and pTi15955 are activated by octopine and AHL, presumably by one or both TraR proteins (TraR^Ti^ and TraR^At^), but that the *traI*^Ti^ gene is more strongly expressed than *traI*^At_1^ (Figure 1B). As such, one explanation for the reduced conjugation frequencies in the Δ*traI*^Ti^ mutant is that although TraI^At_1^ generates similar basal levels of AHL in the absence of octopine induction (Figure 2B and 2C), its expression is less strongly activated by TraR^Ti^ and thus does not create the same level of positive feedback to drive maximal AHL production. We performed plate-based AHL detection assays with the isogenic *traI* mutants harboring the *P_lac_*-*traR*^Ti^ plasmid (Figure 4A). Ectopic expression of *traR*^Ti^ clearly drives strong AHL synthesis in wild type, consistent with positive autoregulation of AHL production by activated TraR. This elevated AHL production was not observed in the Δ*traI*^Ti^ mutant or in the Δ*traI*^Ti^Δ*traI*^At_1^ double mutant, but was clear for the Δ*traI*^At_1^ mutant (Figure 4A). This observation suggested that the weak TraR^Ti^ activation of *traI*^At_1^ expression resulting in less pronounced increases in AHL levels might underlie the diminished conjugation observed in the *traI*^Ti^ mutant. Conversely, we explored how *traR*^At^ impacted AHL production with the isogenic *traI* derivatives expressing *traR*^At^ (Figure 4B). In contrast to *traR*^Ti^, AHL production was enhanced by expression of *traR*^At^ when either *traI* gene was present (Figure 4B).

**Figure 4.**
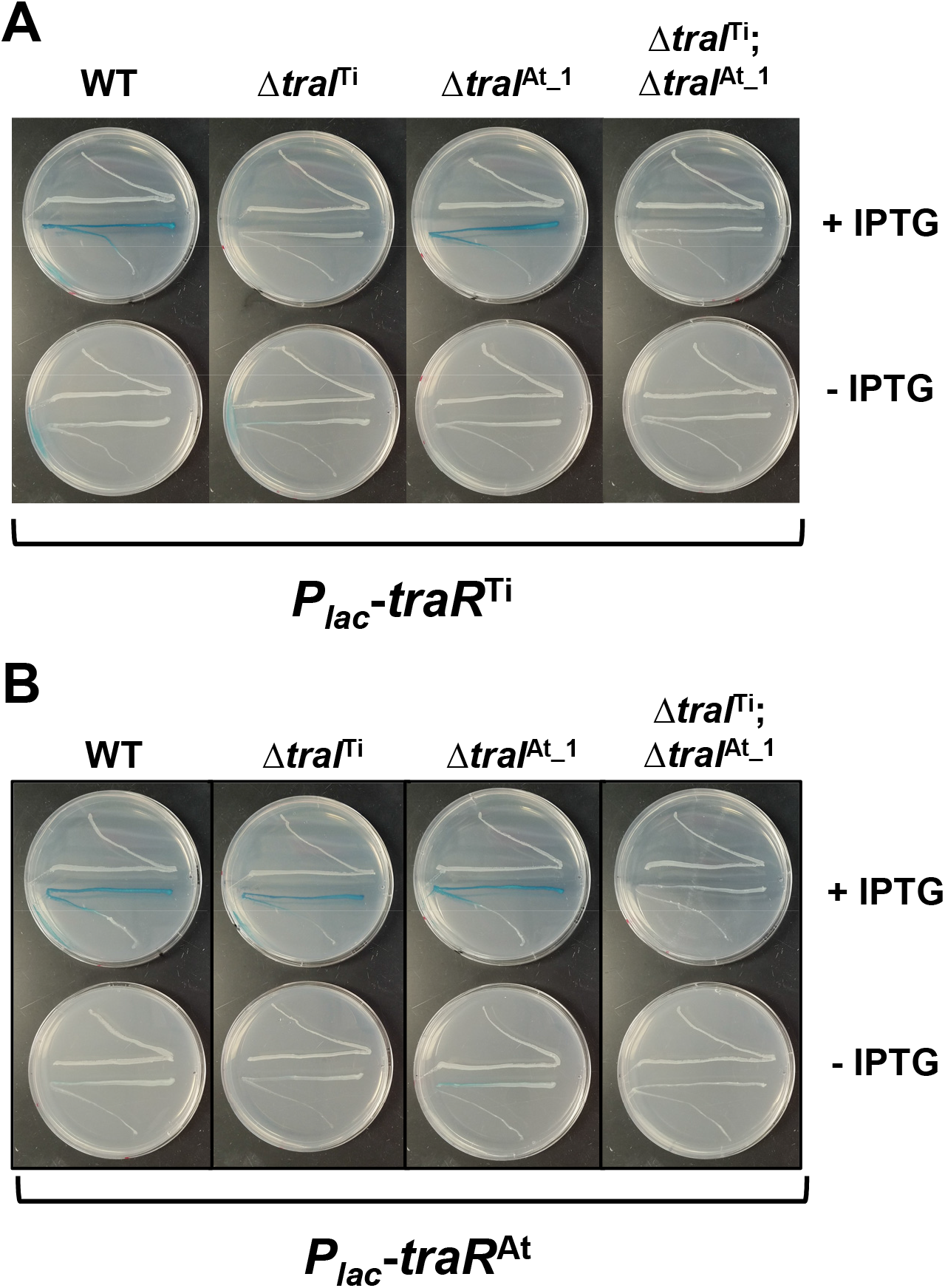
AHL production induced by ectopic expression of *traR*^Ti^ and *traR*^At^ in *A. tumefaciens* 15955. *A. tumefaciens* 15955 expressing a plasmid-borne copy of **(A)***traR*^Ti^ **or (B)***traR*^At^ with or without in-frame deletions in *traI*^Ti^, *traI*^At_1^, or both (top streaks) were used as AHL donors for *A. tumefaciens* NTL4(pCF218)(pCF372) (bottom streaks) on ATGN media supplemented with XGal, with or without 400 μM IPTG (top and bottom plates, respectively). Genotypes of *A. tumefaciens* 15955 AHL donor strain relative to *traI* indicated above.

### Mutational analysis *of traR* genes reveals overlapping control of target promoters

In frame deletions were created in the *traR*^Ti^ (Δ*traR*^Ti^) and *traR*^At^ (Δ*traR*^At^) genes of *A. tumefaciens* 15955 to generate single mutants as well as a double mutant (Δ*traR*^Ti^Δ*traR*^At^). The plasmids carrying *lacZ* fusions to the *traI*^Ti^ and *traI*^At^ genes were independently introduced into these mutants, and β-galatosidase activity ws measured following growth on solid medium under a range of different conditions (Figure 5A and 5B). In wild type *A. tumefaciens* 15955, the *traI*^Ti^*-lacZ* fusion was strongly induced on media with octopine (3.25 mM) with or without the exogenous AHL 3-oxo-C8-HSL (0.1 μM) (Figure 5A). Octopine-dependent induction was abolished in the Δ*traR*^Ti^ mutant, but retained in the Δ*traR*^At^ mutant. Basal level activity was also decreased in the Δ*traR*^Ti^ mutant but a modest increase was observed in the presence of AHL (Figure 5A). In contrast, basal activity from this *lacZ* fusion was slightly increased in the Δ*traR*^At^ mutant. The double mutant lacking both *traR* genes had very low activity that was unaffected by any of the different conditions.

**Figure 5.**
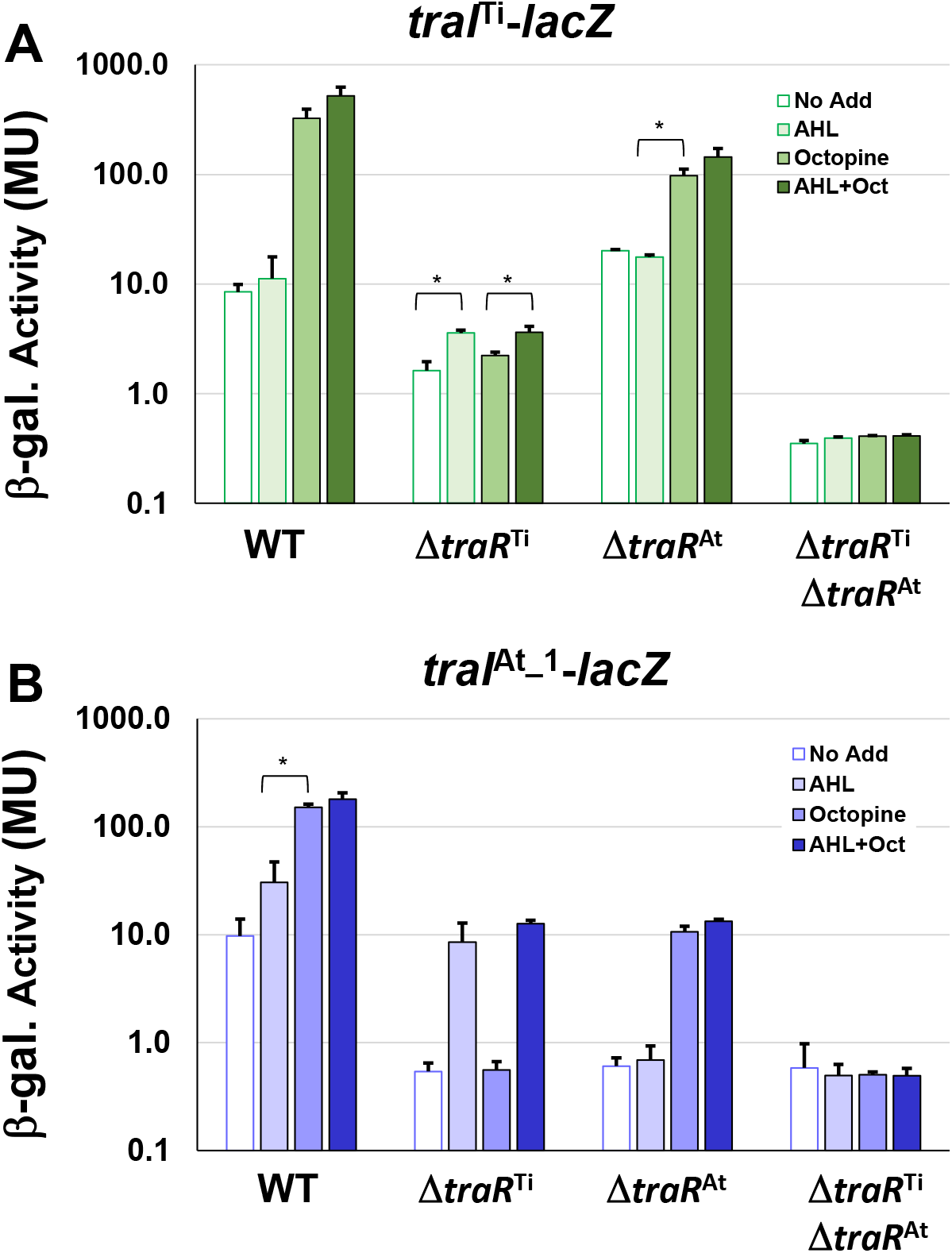
Mutational analysis of *traR* gene with *traI* reporter fusions. WT *A. tumefaciens* 15955 and mutant derivatives (Δ*traR*^Ti^ and Δ*traR*^At^), and (Δ*traR*^Ti^Δ*traR*^At^) carrying a plasmid-borne copy of either *PtraI* from pTi15955 (A, green bars) or from pAt15955 (B, blue bars) translationally fused to *lacZ* were spotted on 0.2 μm cellulose acetate filter discs placed on ATGN on its own (open bars), with 0.1 μM N-3-oxo-C8-HSL (AHL, light-fill bars), with 3.25 mM octopine (Oct., medium-fill bars), and with both (dark-fill bars) Cells were resuspended and promoter activities were calculated as Miller Units (Methods). Bars are standard deviation. All bars that differ by 10-fold or more are significant (p-value <0.05). Bars that differ by less than 10-fold, that are however significantly different (p-value <0.05) are designated with a bracketed asterisk.

Expression of the *traI*^At_1^-*lacZ* fusion showed a different pattern of regulation in the mutant backgrounds. In the wild type background AHL alone was able to significantly increase β-galactosidase activity roughly 3-fold, and addition of octopine with and without exogenous AHL increased this even further (Figure 5B). In the Δ*traR*^Ti^ mutant the overall level of β-galactosidase decreased about 10-fold relative to wild type, but addition of AHL conversely increased the activity 10-fold from that amount. Octopine addition had no effect in the Δ*traR*^Ti^ mutant. The activity imparted from the *traI*^At_1^-*lacZ* fusion also decreased 10-fold in the *traR*^At^ mutant relative to wild type, but in contrast to the *traR*^Ti^ mutant exogenous addition of AHL alone had no effect, whereas octopine or AHL plus octopine both resulted in a 10-fold induction. As with the *traR*^Ti^*-lacZ* fusion, low level activity from the *traI*^At_1^*-lacZ* fusion was unaffected by either octopine or AHL alone or in combination in the Δ*traI*^Ti^Δ*traI*^At_1^ mutant (Figure 5B).

Introduction of compatible plasmids expressing either the *traR*^Ti^ or *traR*^At^ gene from the *P*_lac_ promoter into wild type 15955 and all three different *traR* mutants resulted in strong IPTG-inducible β-galactosidase activity from both the *traI*^Ti^*-lacZ* and *traR*^At^-*lacZ* fusions, in the absence of exogenous AHL or octopine (Figure S5). This is consistent with other *A. tumefaciens* strains where ectopic expression of the *traR* protein, and its positive impact on AHL production, can bypass the need for conjugal opine addition or the effect of exogenous AHL (Fuqua and Winans 1994). Ectopic expression of the *traR*^Ti^ gene however stimulated notably lower β-galactosidase activity from the *traI*^At_1^*-lacZ* fusion than it did for *traI*^Ti^*-lacZ*, and this decreased even further in the Δ*traR* mutant backgrounds, with the lowest activity in the Δ*traR*^Ti^Δ*traR*^At^ mutant. Ectopic expression of *traR*^At^ seemed to have an equivalent impact on both promoters, and this was largely unaffected in the *traR* deletion mutants.

### *In vivo* target gene specificity for TraR^Ti^ and TraR^At^

Our observations on plasmid conjugation, AHL production, and *tra* gene expression suggested that TraR^Ti^ and TraR^At^ might have different specificities for target promoters. Well-studied TraR proteins are known to bind upstream of regulated promoters at conserved *tra* box elements (Fuqua and Winans 1996, Zhu and Winans 1999), sequence motifs that can readily be identified on pTi15955 upstream of *tra* genes. Sequence analysis of the presumptive promoter regions of conjugal transfer genes on pAt15955 identified similar inverted repeat, *tra* box-type elements (Figure S6A and S6B), suggesting direct regulation by one or both TraR proteins. However, the *tra* box consensus motif for pAt15955-encoded genes is slightly different from that of the pTi15955 elements (Figure S6B).

The *traI-lacZ* and *traA-lacZ* fusions from pAt15955 and pTi15955 were introduced into the AHL^−^Ti plasmid-cured *A. tumefaciens* NTL4 strain harboring either the *traR*^Ti^ *or traR*^At^ expression plasmids, that provide IPTG-inducible expression. Very low-β-galactosidase activity was imparted by all of the fusions in the absence of exogenous AHL, even with high level expression of either *traR* gene in the presence of 800 μM IPTG (Figure 6). In the presence of AHL, the *traA*^At^-*lacZ* was strongly activated by *traR*^At^, but about 4-fold more weakly activated by *traR*^Ti^, whereas the difference for activation of the *traA*^Ti^*-lacZ* fusion was less pronounced (Figure 6A, compare blue-filled bars relative to green-filled bars). Activation patterns for the *traI* fusions were similar to the *traA* fusions, though the magnitude of β-galactosidase activity was many fold greater (Figure 6B). The *traI*^At_1^-*lacZ* fusion was again more weakly activated by *traR*^Ti^ than by *traR*^At^ (blue-filled bars), with the activation of the *traI*^Ti^-*lacZ* closer to equivalency for the two regulators (compare green-filled bars). Thus, for both gene fusions derived from pAt15955 the Ti-plasmid encoded *traR* leads to weaker induction, whereas the equivalent gene fusions from pTi15955 are strongly activated by both the TraR^Ti^ and TraR^At^.

**Figure 6:**
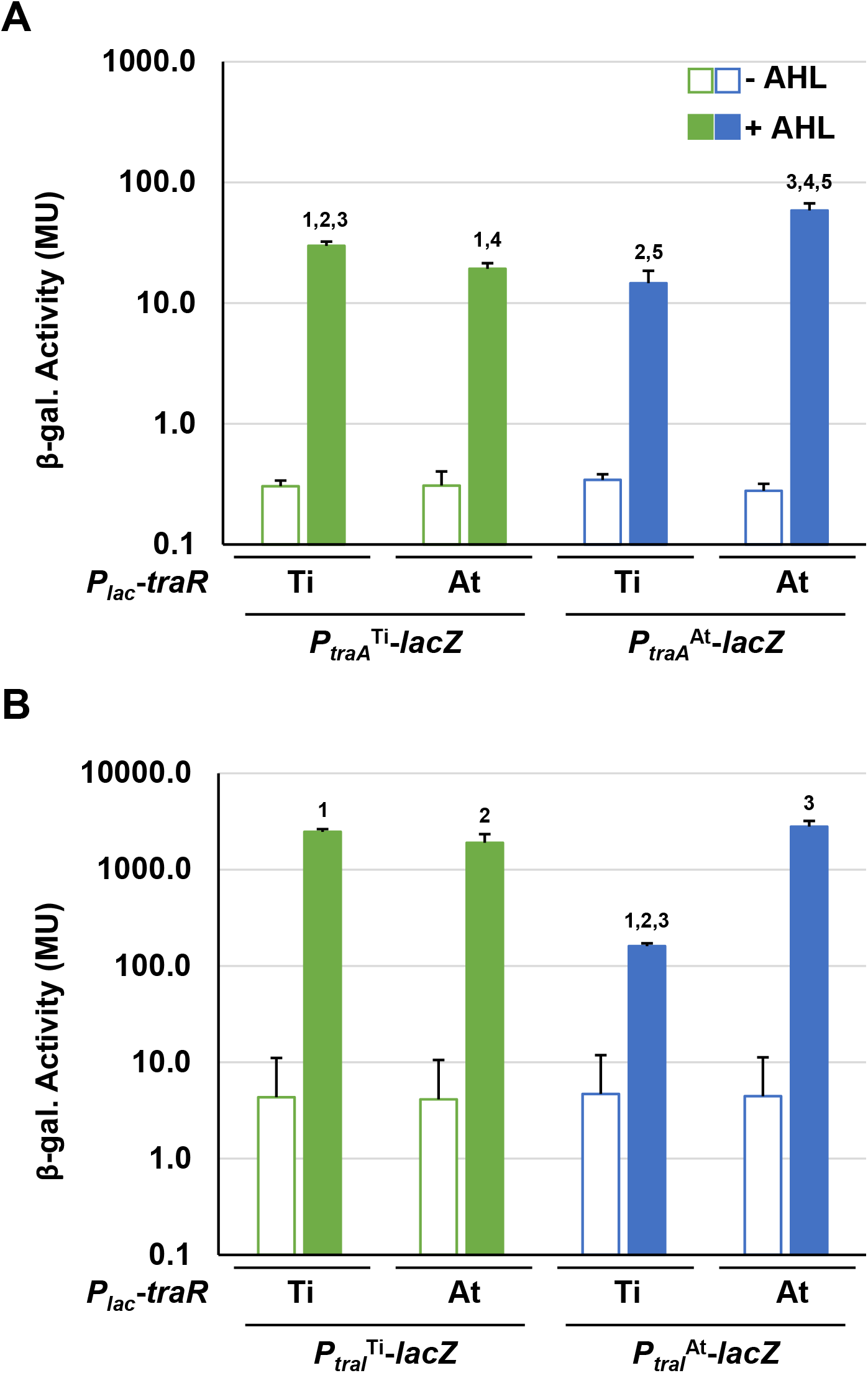
Differential activation of pAt15955 and pTi15955 *P_traA_* and *P_traI_* by *traR*. Plasmid-borne copies of *P_traA_*-*lacZ* or *P_traI_*-*lacZ* from pTi15955 (green) or pAt15955 (blue) and *P_lac_*-*traR* with *traR* genes from either pAt15955 (At) or pTi15955 (Ti) were moved into *A. tumefaciens* NTL4 and β-galactosidase activity (MU) of *P_traA_* (A) or *P_traI_* (B) were determined in the presence or absence of 0.1 μM 3-oxo-C8-HSL (+ AHL and – AHL, respectively). Bars with the same numbers possess significant difference in averages (p-value <0.05).

## DISCUSSION

Members of the Family *Rhizobiacea*e often harbor multiple *repABC*-based secondary replicons that are compatible with each other and are replicated and segregated faithfully during the cell cycle (Castillo-Ramirez et al. 2009, Cevallos et al. 2008). Unlike secondary chromosomes and chromids, megaplasmids and plasmids within this group often encode conjugation machinery and undergo copy number fluctuations tied with functions such as virulence, symbiosis, and horizontal transfer. The Ti plasmid from *A. tumefaciens* is one of the most well-studied *repABC*-controlled replicons, where virulence and horizontal transfer are tightly controlled by environmental signals. In addition to proper coordination with the cell cycle, tight regulation of copy number and horizontal gene transfer of large secondary replicons serves to limit the costs associated with DNA replication and horizontal transfer.

The opine-dependent quorum sensing system is well characterized in *Agrobacterium* species harboring the wide host range octopine-type and nopaline-type Ti plasmids, with prototypes pTiR10 and pTiC58, respectively (Gordon and Christie 2014). Based on extensive genomic sequencing the *Agrobacterium* genus has been phylogenetically subdivided into over 10 different genomospecies (Lassalle, et al. 2011), with pTiR10 found in Genomospecies 4 and pTiC58 found in Genomospecies 8. In these systems there is a single quorum sensing system comprised of the TraR transcription factor, the TraI AHL synthase, and the TraM antiactivator, which together regulate Ti plasmid conjugation and copy number. However, in several members of the *Agrobacterium* G1 genomospecies with octopine-type Ti plasmids, such as *A. tumefaciens* 15955, *A. tumefaciens* A6, and *A. tumefaciens* Ach5, the At plasmid carries a second TraR-TraI-TraM quorum sensing system, that is closely related to the Ti-encoded regulators. We have demonstrated that this second At-encoded system controls conjugative transfer through regulation of *tra* genes on the At plasmid, but that it can also have a profound impact on Ti plasmid functions. Likewise, the Ti-encoded system can impact At plasmid conjugative transfer, most notably imparting octopine-inducible control over pAt15955 conjugation (Barton, et al. 2019). These co-resident quorum sensing systems interdigitate at multiple levels to regulate horizontal transfer of both plasmids (Fig. 7). Both TraI gene products are highly similar to each other, and drive AHL synthesis. Although the chemical identity of the AHL is not formally shown here for *A. tumefaciens* 15955, is has ben reported for the closely related *A. tumefaciens* A6 that both the Ti and At *traI* genes specify production of 3-oxo-C8-HSL (Wang et al. 2014). The TraI proteins between A6 and 15955 show 100% amino acid identity. Furthermore, we show here that either *traI* gene can function to facilitate conjugation of both plasmids, but loss of both *traI* genes together abolishes conjugation of each plasmid. Therefore, we predict that the two TraI proteins in *A. tumefaciens* 15955 primarily contribute to the same pool of 3-oxo-C8-HSL (Figure 7). This pool of AHL is required for TraR-mediated activation of *tra* genes and plasmid conjugation, through protein-ligand interactions with their amino-terminal ligand binding domains.

**Figure 7:**
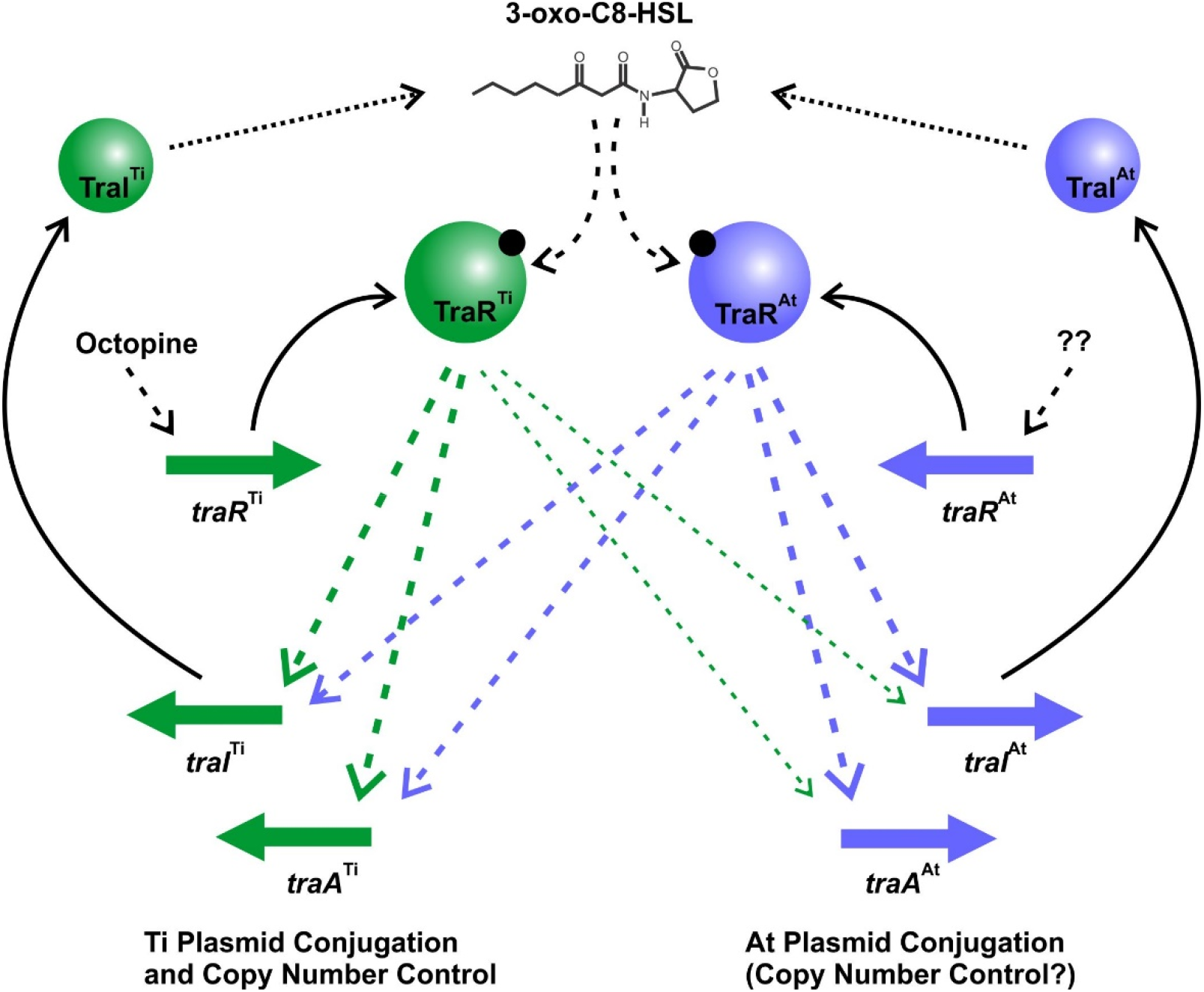
Model of Dual Quorum Sensing Systems in *A. tumefaciens* 15955. Green shaded symbols representing proteins and genes from pTi15955, and blue shaded symbols represent proteins and genes from pAt15955. Solid arrows indicate translation of specific genes, dotted arrows represent enzymatic synthesis of AHL, and dashed arrows are regulatory impacts of each system. The thinner arrows from TraR^Ti^ to the pAt15955 target genes reflect the observed weaker effect of this protein on their expression.

In the presence of inducing levels of AHL the TraR proteins can bind to DNA and activate expression of target promoters, and stimulate their conjugative transfer. The presence of octopine, naturally produced by plant tissue transformed with T-DNA encoding the octopine synthase genes, has been shown to activate expression of the Ti plasmid *traR* gene in other octopine-type plasmids (Fuqua and Winans 1994). The operon structure upstream of the pTi15955 *traR* is identical to that on these better studied octopine-type Ti plasmids (Fig. S1B). We posit that octopine increases expression of *traR*^Ti^ on pTi15955 as well, and that this increase stimulates expression of the *traI*^Ti^ gene, which in turn elevates AHL levels, creating a positive feedback loop as observed for many LuxR-LuxI-type systems (Fuqua and Greenberg 2002). Ectopically increasing expression of *traR*^Ti^ bypasses the need for octopine. However, *traR*^Ti^ only weakly stimulates the *traI* gene on the At plasmid, which in turn limits its impact on AHL synthesis, thereby explaining the weaker stimulation of both Ti plasmid and At plasmid conjugation in the Δ*traI*^Ti^ mutant (Figure 3). The *traR*^At^ gene product appears to activate expression of both the *traI*^At_1^ and *traI*^Ti^ with rough equivalence. However, there is no evidence that its expression is affected by octopine, and consistent with this, in the absence of the *traR*^Ti^ gene octopine has no impact on the *tra* gene expression or conjugation of either plasmid. We hypothesize that under laboratory conditions, *traR*^At^ expression is low, and insufficient to stimulate significant positive feedback on AHL production, and thus pTi15955 conjugative transfer is extremely infrequent or undetectable in the absence of octopine. The basal expression of the *traI*^Ti^ gene is in fact modestly elevated in the Δ*traR*^At^ mutant raising the possibility that TraR^At^ plays a mildly inhibitory role in the absence of octopine. In contrast to pTi15955, low level conjugation of pAt15955 is detectable in the absence of octopine (Barton et al. 2019). The observation that even this minimal conjugation is largely abolished in the Δ*traI*^Ti^Δ*traI*^At^ double mutants suggests that quorum sensing is functioning to regulate the process even at its lowest levels. It also is plausible that expression of the *traR*^At^ gene is regulated by a condition that we have not yet identified such as has been observed in other rhizobia, and under certain conditions might elevate pAt15955 and thereby indirectly pTi15955 conjugation (Banuelos-Vazquez et al. 2020).

The molecular mechanisms by which the two TraR proteins function in the same cell remains an area for future studies. There are abundant examples of multiple LuxR-type regulators functioning in a single bacterial species, often in some form of hierarchal network. For example, in *Pseudomonas aeruginosa* there are three distinct LuxR-type regulators, LasR, RhlR, and QscR with overlapping regulons (Ding et al. 2018). In other Alphaproteobacteria multiple LuxR proteins are also common. For example, in the marine bacterium *Ruegeria* sp. KLH11 the SsaR LuxR-type protein activates motility in response to long chain AHLs generated by the SsaI AHL synthase, and also regulates the activity of a second LuxR-type protein SsbR and two AHL synthases SsbI and SscI (Zan et al. 2015, Zan et al. 2012). Multiple, co-resident LuxR-LuxI-type proteins are well documented in the rhizobia, and their regulatory interactions are notably complex and convoluted. *Rhizobium leguminosarum* biovar *viciae* has a cascade of multiple LuxR and LuxI-type proteins including several that control plasmid conjugation (Lithgow et al. 2002, Cervantes, et al. 2019). The biocontrol agent known as *Rhizobium rhizogenes* K84 encodes two TraR homologues (TraR_acc_ and TraR_noc_) on the single pAtK84b plasmid that combine to control conjugative transfer of this plasmid (Oger et al. 1998). Remarkably, in K84 expression of each *traR* gene is regulated by a different conjugal opine, nopaline and agrocinopine A/B, and consequently both opines independently stimulate pAtK84b conjugation through a single conjugative machinery. Recently, two of the *repABC* plasmids of *Sinorhizobium fredii* GR64, pSfr64a and pSfr64b have been reported to encode two separate TraR-TraM-TraI quorum sensing systems, in addition to a quorum sensing regulatory system encoded by the chromosome, NgrR-NgrI (Cervantes, et al. 2019). All three of these quorum sensing systems impact each other and contribute to control of pSfr64a and pSfr64b conjugation. Similar to *S. fredii* GR64, *A. tumefaciens* 15955 and its immediate relatives have dual, highly conserved TraR-TraI-TraM systems on two separate plasmids that each encode their own conjugative transfer machinery. Although there is no chromosomal LuxR-LuxI system in *A. tumefaciens* 15955, our findings reveal that the plasmid encoded quorum sensing systems are clearly interdependent. Dimerization is known to be a requirement for DNA binding among diverse LuxR homologues including TraR. Given the similarity of the TraR^Ti^ and TraR^At^ proteins (63.2% identity), heterodimer formation is plausible, potentially altering the activity of each TraR protein. Interestingly, *A. tumefaciens* 15955 is known to encode a truncated copy of the *traR*^Ti^ gene, named TrlR that is 88% identical, but ablated for the DNA binding domain, expression of which is inducible by the non-conjugative opine mannopine (Oger, et al. 1998, Zhu, et al. 1998). TrlR is a dominant negative homologue of TraR, that forms an inactive heterodimer with TraR^Ti^ and prevents DNA binding (Chai et al. 2001). In the absence of mannopine, TrlR is not expressed. Our genetic analysis revealed that mutation of *traR*^At^ modestly elevated expression of the *traI*^Ti^*-lacZ* fusion in response to octopine, suggesting that TraR^At^ may also have a mild dominant negative effect on TraR^Ti^.

Another component of the quorum sensing systems commonly encoded on Ti plasmids is the TraM anti-activator (Fuqua, et al. 1995, Hwang, et al. 1995). TraM forms a multimeric complex with TraR, abrogating its activity and targeting it for proteolysis (Chen et al. 2007, Costa et al. 2012, Swiderska et al. 2001). *A. tumefaciens* 15955 encodes TraM proteins on pAt15955 and pTi15955, that are 65% identical. Genetic and biochemical analysis of a second TraM homologue from the close relative *A. tumefaciens* A6 demonstrated that it can form a complex with TraR encoded by the Ti plasmid and adopts the same three-dimensional structure as TraM^Ti^ (Chen et al. 2006, Wang, et al. 2014). The two TraM proteins seemed to be functionally redundant in *A. tumefaciens* A6, and mutation of both genes was required to disregulate the Ti plasmid TraR. Subsequent studies prior to genome sequencing also identified *traR* and *traI* genes linked to this second *traM* gene, all of which we now recognize to be carried on the pAtA6 plasmid (Wang, et al. 2014). Although we did not examine the role for the *traM*^Ti^ and *traM*^At^ in the current study on *A. tumefaciens* 15955, it seems very likely that they play similar roles as those in A6. As such, the orchestration between the pTi15955 and pAt15955 quorum sensing systems to control plasmid transfer, very likely also includes these TraM anti-activators and their ability to inhibit their cognate and non-cognate TraR proteins.

We observed that TraR^Ti^ only weakly activated the pAt15955 gene fusions tested, while it strongly affected the homologous pTi15955 gene fusions. In contrast, TraR^At^ was largely equivalent in activation of both pTi15955 and pAt15955 targets, suggesting that it has a more permissive DNA binding specificity. Comparatively, the *tra* box elements identified upstream of these and other potential target genes, are similar 18 bp inverted repeats, but there are several differences at specific positions, that may distinguish the pTi15955 targets from the pAt15955 sequences. Amino acid sequence differences between TraR^Ti^ and TraR^At^ within the C-terminal DNA-binding domain are likely to impart the DNA binding site specificity. Although the proteins are quite similar overall in the DNA binding domain (69% identity), there are several differences including a conserved isoleucine residue (I194) in TraR^Ti^ (either an I or V in most LuxR-type proteins, (Whitehead, et al. 2001)), that is an alanine (A194) in TraR^At^ (Figure S6). The octopine-type *A. tumefaciens* R10 TraR protein which is encoded on pTiR10, was structurally characterized in association with the 3-oxo-C8-HSL ligand and bound to a synthetic *tra* box oligonucleotide (Zhang et al. 2002). Two of the primary contacts with DNA residues within the *tra* box were Arg206 (R206) and (R210) which are conserved in the 15955 TraR^Ti^ protein (Fig. S4, asterisks). However, these positions are divergent in 15955 TraR^At^ and are Lys206 and Glu210 (Fig. S4). Thus, it seems the potential mechanism of *tra* box recognition may be different between these two regulators. Our findings suggest that sequence divergence between the two proteins imparts a less restrictive DNA binding site specificity to TraR^At^ than to TraR^Ti^. However, testing the binding of each TraR paralogue to pAt15955 and pTi15955 promoters *in vitro* would provide an effective way to directly evaluate their specificity.

## Supporting information

Manuscript file plus figures

## Acknowledgements

This work was supported by National Institutes of Health Grant (R01 GM092660 to C.F. and James D. Bever), the Indiana University Faculty Research Support Program (FRSP) (to C.F.), the National Science Foundation (NSF) Graduate Research Fellowship Program (GGVP004842 to P.A.N.-O.), and the Kansas NSF EPSCoR First Award Program (OIA-1656006 to T.G.P.). We would also like to thank Isam Madi and Reilly Jensen for help with strain construction.

