## Supplementary material for "Co-Dependent and Interdigitated: Dual Quorum Sensing Systems Regulate Conjugative Transfer of the Ti Plasmid and the At Megaplasmid in *Agrobacterium tumefaciens* 15955": Manuscript file plus figures

Supplementary Tables (S1-S3), Supplementary Figure Legends, Supplementary References, Supplementary Figures (S1-S6)

### SUPPLEMENTAL TABLES

#### Table S1: Strains used in this study

| Strain | Genotype/markers | Notes | Reference |
|---|---|---|---|
| ***E. coli*** | | | |
| DH5α/λpir | λpir ; cloning strain | | (Chiang and Rubin 2002) |
| S17-1/λpir | λpir ; $Tra^{+}$ , cloning host | | (Kalogeraki and Winans 1997) |
| ***A. tumefaciens*** | | | |
| C58-ERM52 | $Sp^{R}$ | Plasmidless (pAt-, pTi-) Conjugation recipient | (Morton et al. 2014) |
| NTL4 | C58 Δ*tetRA* | pTi-cured derivative of *A. tumefaciens*, $AHL^{-}$ | (Luo et al. 2001) |
| 15955-KT1 | Δ*traI*$^{Ti}$ | | This study |
| 15955-KT2 | Δ*traI*$^{At\_1}$ | | This study |
| 15955-KT3 | Δ*traI*$^{Ti}$; Δ*traI*$^{At\_1}$ | | This study |
| 15955-IB123 | pTi15955::*gusA*$Gm^{R}$ | | Barton et al. 2019 |
| 15955-IB125 | pAt15955::$Km^{R}$ | | Barton et al. 2019 |
| 15955-IB137 | pAt15955::$Km^{R}$; Δ*traI*$^{Ti}$ | KT1, marked on pAt15955 with $Km^{R}$ | This study |
| 15955-IB138 | pAt15955::$Km^{R}$; Δ*traI*$^{At\_1}$ | KT2, marked on pAt15955 with $Km^{R}$ | This study |
| 15955-IB139 | pAt15955::$Km^{R}$; Δ*traI*$^{Ti}$; Δ*traI*$^{At\_1}$ | KT3, marked on pAt15955 with $Km^{R}$ | This study |
| 15955-IB148 | pTi15955::*gusA*$Gm^{R}$; Δ*traI*$^{Ti}$ | KT1, marked on pTi15955 with *gusA* and $Gm^{R}$ | This study |
| 15955-IB149 | pTi15955::*gusA*$Gm^{R}$; Δ*traI*$^{At\_1}$ | KT2, marked on pTi15955 with *gusA* and $Gm^{R}$ | This study |
| 15955-IB150 | pTi15955::*gusA*$Gm^{R}$; Δ*traI*$^{Ti}$; Δ*traI*$^{At\_1}$ | KT3, marked on pTi15955 with *gusA* and $Gm^{R}$ | This study |
| 15955-PAG1 | Δ*traR*$^{Ti}$ | | This study |
| 15955-IAM1 | Δ*traR*$^{At}$ | | This study |
| 15955-PAG9 | Δ*traR*$^{Ti}$Δ*traR*$^{At}$ | | This study |

**Table S2: Plasmids used in this study**

| Plasmid | Features | Reference |
|---|---|---|
| pIB302 | pSRKGm $P_{lac}$-*traI*$^{At\_1}$ | This study |
| pIB303 | pSRKGm $P_{lac}$-*traI*$^{Ti}$ | This study |
| pIB305 | pSRKGm $P_{lac}$-*traI*$^{At\_2}$ | This study |
| pIB306 | pSRKGm $P_{lac}$-*traR*$^{At}$ | This study |
| pIB307 | pSRKGm $P_{lac}$-*traR*$^{Ti}$ | This study |
| pIB308 | pSRKKm $P_{lac}$-*traR*$^{At}$ | This study |
| pIB309 | pSRKKm $P_{lac}$-*traR*$^{Ti}$ | This study |
| pIB310 | pRA301 $P_{traA}$$^{At}$-*lacZ* | This study |
| pIB311 | pRA301 $P_{traI}$$^{At}$-*lacZ* | This study |
| pIB312 | pRA301 $P_{traA}$$^{Ti}$-*lacZ* | This study |
| pIB313 | pRA301 $P_{traI}$$^{Ti}$-*lacZ* | This study |

**Table S3: Primers used in this study***

**Table S3: Primers used in this study***

| Primer | Sequence | Description |
|---|---|---|
| IBP156 | aggcatatgttggagcggac | pAt15955 *traI*_2 (truncated) 5' w/NdeI for expression in pSRK |
| IBP157 | aggactagttcatccatctttgc | pAt15955 *traI*_2 (truncated) 3' w/SpeI for expression in pSRK |
| IBP158 | aggcatatgcgaatcttgac | pAt15955 *traI*_2 (modified) 5' w/NdeI for expression in pSRK |
| IBP159 | ggaactagttcatccatctttgc | pAt15955 *traI*_2 (modified) 3' w/SpeI for expression in pSRK |
| IBP160 | ggacatatgcaacattggct | pAt15955 *traR* 5' w/NdeI for expression in pSRK |
| IBP161 | aggactagttcagatcagcccgg | pAt15955 *traR* 3' w/SpeI for expression in pSRK |
| IBP162 | ggacatatgcagcactggct | pTi15955 *traR* 5' w/NdeI for expression in pSRK |
| IBP163 | aggactagttcagatgagtttcc | pTi15955 *traR* 3' w/SpeI for expression in pSRK |

| | | |
|---|---|---|
| IBP190 | aggcatatgcggatcctgaccgttcc | pAt15955 *traI*_1 5' w/NdeI for expression in pSRK |
| IBP191 | aggactagttcacgccgcgctcctcgccg | pAt15955 *traI*_1 3' w/SpeI for expression in pSRK |
| IBP192 | aggcatatgctgattctgaccgtctc | pTi15955 *traI* 5' w/NdeI for expression in pSRK |
| IBP193 | aggactagttcacgccgcactcctcaacg | pTi15955 *traI* 3' w/SpeI for expression in pSRK |
| IBP210 | gaattcgagctcggtaccttttgcgcgcatctgatgtcatcg | pRA301 (SphI/KpnI) *lacZ* fusion, *traA*$^{pAt}$ 5’ |
| IBP212 | gaattcgagctcggtaccagcgttttgtgcgaagtgg | pRA301 (SphI/KpnI) *lacZ* fusion, *traI*$^{pTi}$ 5’ |
| IBP213 | ggagcaagcttgcatgccatgttgatctccgcaacaagaaacga | pRA301 (SphI/KpnI) *lacZ* fusion, *traI*$^{pTi}$ 3’ |
| IBP216 | gaattcgagctcggtaccgctcgctaccggtccggct | pRA301 (SphI/KpnI) *lacZ* fusion, *traA*$^{pTi}$ 5’ |
| IBP217 | ggagcaagcttgcatgccacggcgaagtgcgctcccgg | pRA301 (SphI/KpnI) *lacZ* fusion, *traA*$^{pTi}$ 3’ |
| IBP218 | ggagcaagcttgcatgccacggcgaagagcgctcccgg | pRA301 (SphI/KpnI) *lacZ* fusion, *traA*$^{pAt}$ 3’ |
| IBP219 | gaattcgagctcggtaccccgatttcgccttgatcggg | pRA301 (SphI/KpnI) *lacZ* fusion, *traI*$^{pAt}$ 5’ |
| IBP220 | ggagcaagcttgcatgccatatctttctccgctt | pRA301 (SphI/KpnI) *lacZ* fusion, *traI*$^{pAt}$ 3’ |

Note:

*All other primers used in strain creation or diagnostics are available upon request.

### SUPPLEMENTARY FIGURE LEGENDS

**Figure S1. Comparative gene organization within pAt and pTi from *A. tumefaciens* 15955 and *A. tumefaciens* C58. (A)** Whole replicon gene cluster locations of pAt15955, pAtC58, pTi15955, and pTiC58 from *A. tumefaciens* 15955 and *A. tumefaciens* C58 (also known as *A. fabrum*). Light purple bar labeled pAt15955Δ270 indicates the segment precisely deleted upon curing of pTi15955 previously described (Barton et al. 2019) **(B)** Comparison of *traM-traR* gene neighborhoods for pAt15955, pTi15955 and pTiC58. Gene names and unannotated gene numbers provided from NCBI annotation of the *A. tumefaciens* 15955 and *A. tumefaciens* C58 genome sequences. **(C)** Comparison of *traI* gene neighborhoods for pAt15955, pTi15955 and pTiC58. $\mathrm{TraI}^{\mathrm{At\_2}}$ is included in crosshatching and with a solid outline, but was found to not produce a detectable AHL. Solid black vertical lines on gene maps mark locations of *tra* box elements.

**Figure S2. Clustal omega alignment of TraI sequences from pTi15955 and pAt15955.** Conserved residues within the AHL synthase family are highlighted in red (Churchill and Herman 2008).

**Figure S3. Conjugation of pAt15955 is restored in *A. tumefaciens* 15955 Δ*traI*$^{\mathrm{Ti}}$ Δ*traI*$^{\mathrm{At\_1}}$ upon exogenous addition of 3-oxo-C8-HSL.** *A. tumefaciens* 15955 Δ*traI*$^{\mathrm{Ti}}$ Δ*traI*$^{\mathrm{At\_1}}$ harboring a plasmid-borne copy of $P_{lac}$-*traR* from either pAt15955 (purple) or pTi15955 (green) were mixed 1:1 with a plasmidless recipient (ERM52) and spotted onto ATGN media containing 400 µM IPTG and varying amounts of 3-oxo-C8-HSL (X axis). After 24 hours, conjugation frequencies (Y axis) were calculated as transconjugants per output donor (see Materials and Methods).

**Figure S4. Clustal omega alignment of TraR sequences from pTi15955 and pAt15955.** Conserved residues within the AHL coordination and DNA-binding domains are highlighted in orange or red, respectively. Arrow indicates single amino acid difference within the DNA-binding domain of pAt and pTi TraR, and asterisks mark residues R/K206 and R/E210 in each protein (White and Winans 2007).

**Figure S5. Ectopic expression of either *traR* gene in *traR* null mutants increases expression of *traI* targets.** *A. tumefaciens* 15955 and mutant derivatives ($\Delta$*traR*$^{Ti}$, $\Delta$*traR*$^{At}$, and $\Delta$*traR*$^{Ti}$$\Delta$*traR*$^{At}$) carrying a plasmid-borne copy of *traR* either from pTi15955 (solid bars) or from pAt15955 (striped bars) and *traI-lacZ* fusions from either pTi15955 (green) or pAt15955 (blue) plasmid were spotted on 0.2 µm cellulose acetate filter discs placed on ATGN alone (light fill) or with 400 µM IPTG (dark fill) to induce expression of the *traR* gene, and incubated at 28°C for 48 h. Cells were resuspended and promoter activities were calculated as Miller Units (Methods). Bars are standard deviation. All bars that differ by 10-fold or more are significant (p-value <0.05). Bars that differ by less than 10-fold, that are however significantly different from the corresponding wild type background (p-value <0.05) are designated with an asterisk or double asterisk.

**Figure S6. Alignment of subset of putative *tra* boxes from pAt and pTi in *A. tumefaciens* 15955 and motif analysis.** (A) Inverted repeats of putative *tra* box elements are indicated by black arrows. Approximate -35 and -10 elements and +1 sites are indicated by gray boxes or bolded, respectively. Bolded residues are putative transcriptional start sites. (B) Putative *tra* boxes from pAt15955 or pTi15955 (Figure S10A) were used to create a sequence motif with WebLogo (Crooks et al. 2004).

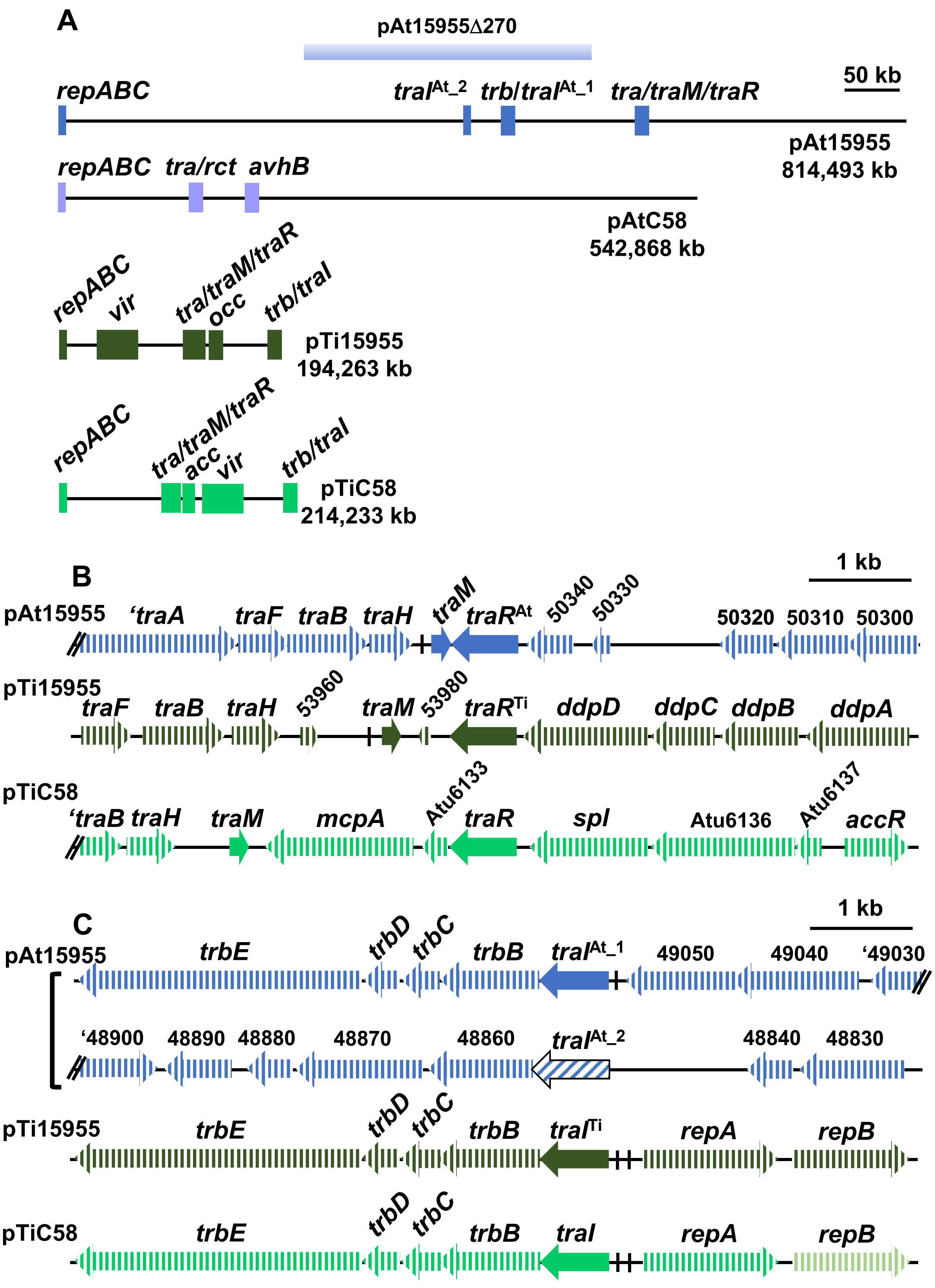


Barton et al.; Figure S1

```
TraI^Ti     MLILTVSPDQYQHQNSYLKQMHRLRAEVFGNRLKWDVAIEDGGERDQYDELSPTYILATF   60
TraI^At_1   MRILTVPPDQHAHHKDELAQMHRLREMVFRGRLEWDVTVTETGEFDEYDQFDPTYVLAVT   60
TraI^At_2   MRILTVPRDQYTDQSDELAQMHRLRAIGFRDRLERGVTTSESGEFDQRDQLDPTNIRAVA   60
            * ****  **: .:.. * ******   * .**: .*:  : ** *: *::.** : *.

TraI^Ti     GGQRVVGCARLLPASGPTMLERTFPQLLATGSLSATTAMIESSRFCVDTTLPTGRAGRQL   120
TraI^At_1   DYGRVVGCARLLPAVGPTMLELTFPQLLRDGSLNATSAMIESSRFCVDTTLPAGRGGGQL   120
TraI^At_2   NDNRVVVCARRLPAVGRTMLERTFPQLLASGAFNAIYRMIESFRVCIDTSFP-GRGGGQF   119
            .  *** *** *** * **** ******  *::.*   **** *.*:**::* **.* *:

TraI^Ti     HLATLTMFAGIIEWSMANGYDEIVTATDLRFERILKRAGWPMTRLGEPVAIGNTVAVAGH   180
TraI^At_1   HLATLTMFAGIIEWSMANGYDKIVTATDLRFERILNRAGWPMARLGEPVAIDNTVAIAGT   180
TraI^At_2   HLTALTVFAFIIELSMANGYDQIVTAKDLDFECTRNRPRWPITRPGEPVEIDNTIAISGT   179
            **::**:** *** *******:****.** **   :*  **::* **** *.**:*::*

TraI^Ti     LPADRKSFERVCPPGYRSIIADDNGRPLRSAA--------------------------   212
TraI^At_1   LPADQESFEQVRRPNYRSTLSYGDNHSARSAA--------------------------   212
TraI^At_2   LPTDQASFEQVCPMGYRSITSGNGVDPAGAKTSAAVPPRMTNLKARRNSAQVRTCKDG   237
            **:*: ***:*   .***  : ..     : :
```

**Barton et al.**
**Figure S2**

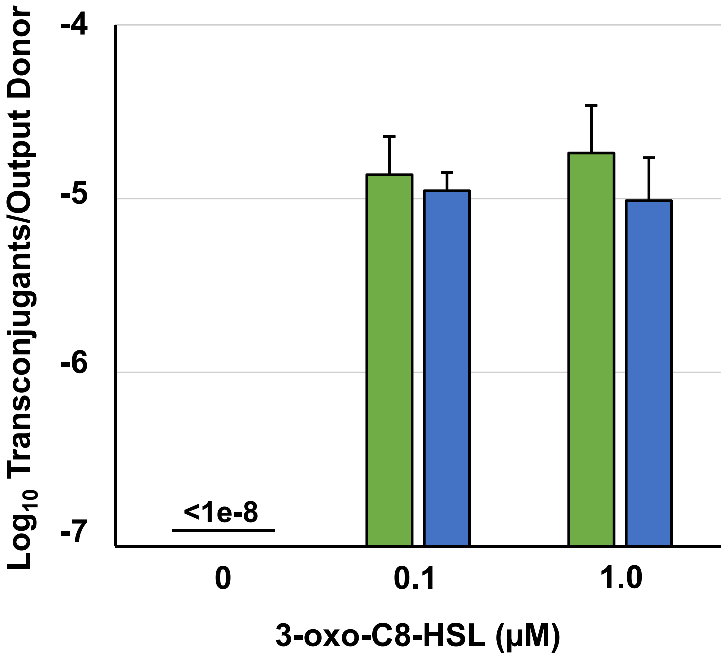


**Barton et al.**
**Figure S3**

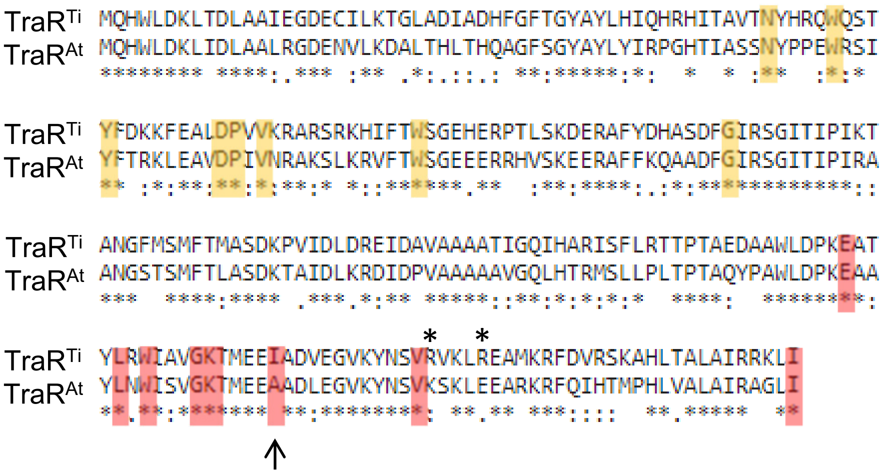


**Barton et al.**
**Figure S4**

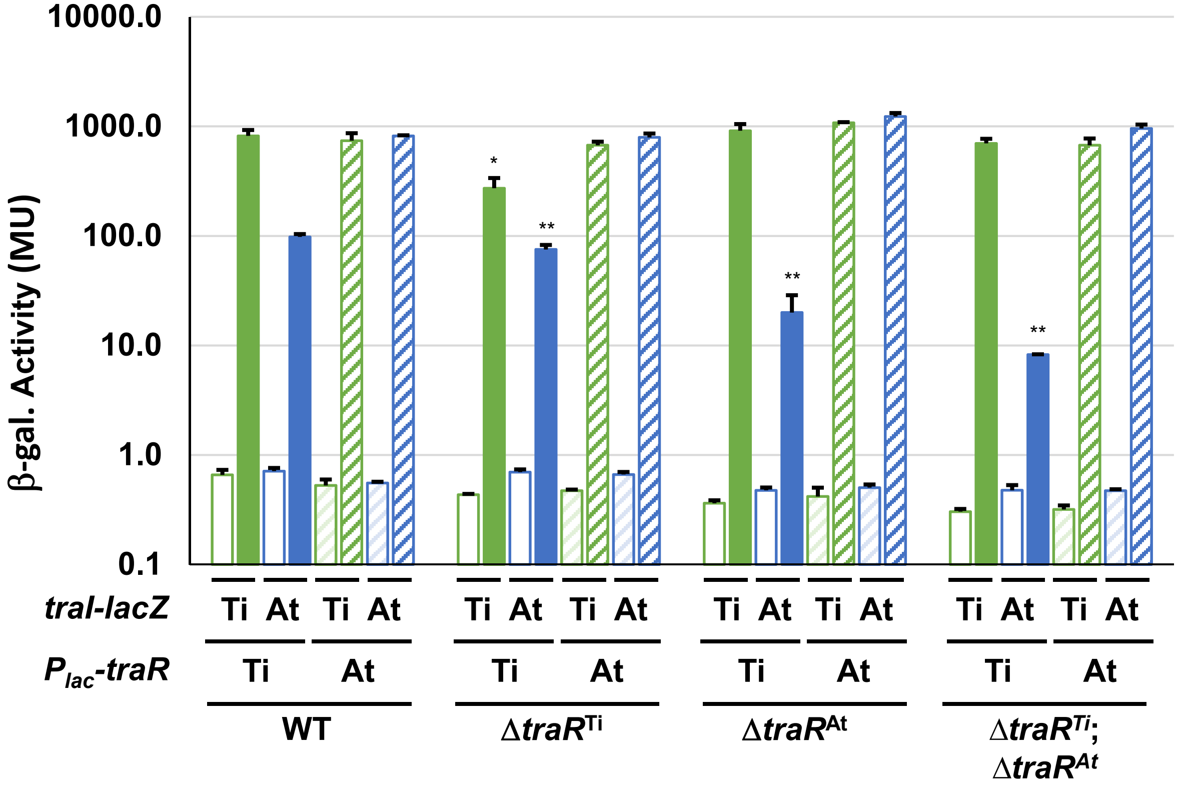


**Barton et al.**
**Figure S5**

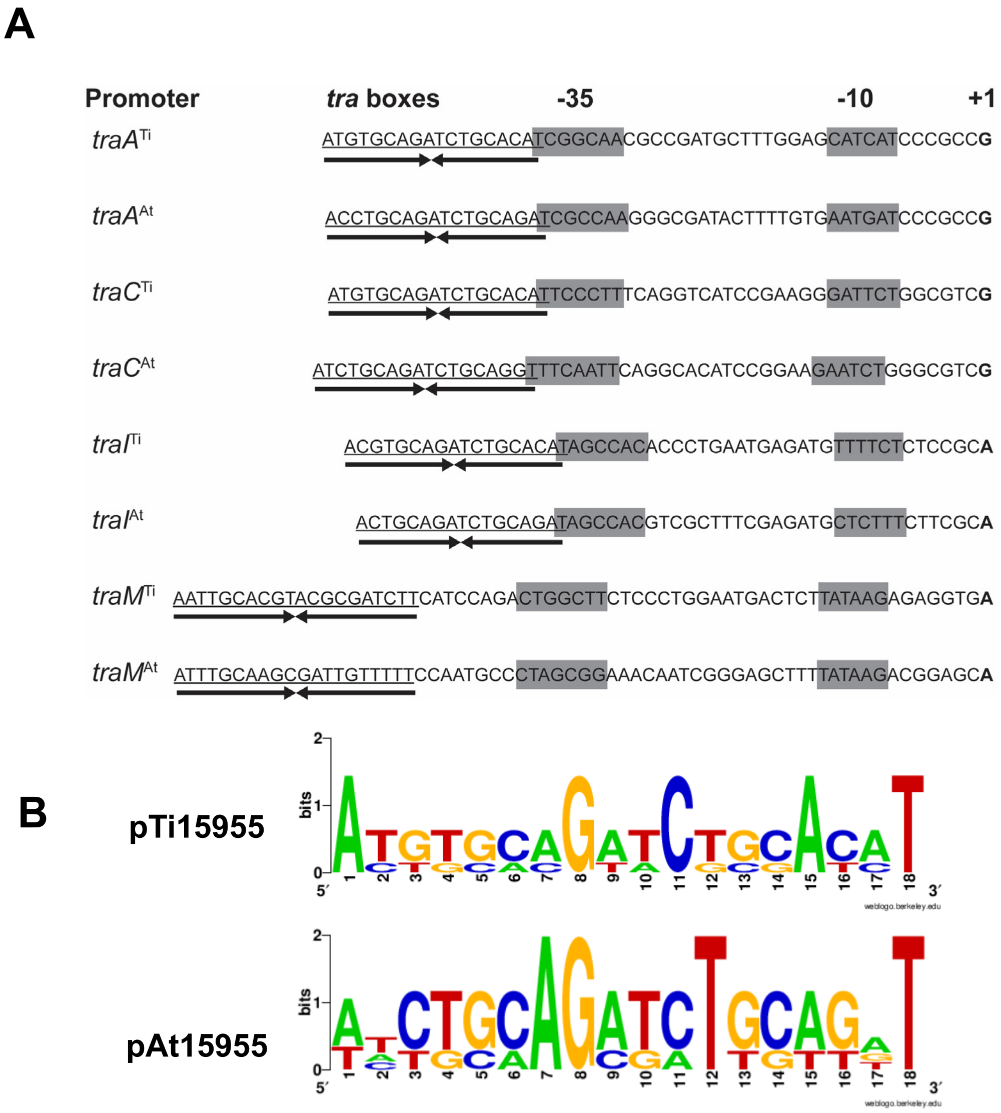


**Barton et al.**
**Figure S6**
